# Bayesian Node Dating based on Probabilities of Fossil Sampling Supports Trans-Atlantic Dispersal of Cichlid Fishes

**DOI:** 10.1101/038455

**Authors:** Michael Matschiner, Zuzana Musilová, Julia M. I. Barth, Zuzana Starostová, Walter Salzburger, Mike Steel, Remco Bouckaert

**Affiliations:** Centre for Ecological and Evolutionary Synthesis (CEES), Department of Biosciences, University of Oslo, Oslo, Norway; Zoological Institute, University of Basel, Basel, Switzerland; Department of Zoology, Faculty of Science, Charles University in Prague, Prague, Czech Republic; Department of Mathematics and Statistics, University of Canterbury, Christchurch, New Zealand; Department of Computer Science, University of Auckland, Auckland, New Zealand; Computational Evolution Group, University of Auckland, Auckland, New Zealand

## Abstract

Divergence-time estimation based on molecular phylogenies and the fossil record has provided insights into fundamental questions of evolutionary biology. In Bayesian node dating, phylogenies are commonly time calibrated through the specification of calibration densities on nodes representing clades with known fossil occurrences. Unfortunately, the optimal shape of these calibration densities is usually unknown and they are therefore often chosen arbitrarily, which directly impacts the reliability of the resulting age estimates. As possible solutions to this problem, two non-exclusive alternative approaches have recently been developed, the “fossilized birth-death” model and “total-evidence dating”. While these approaches have been shown to perform well under certain conditions, they require including all (or a random subset) of the fossils of each clade in the analysis, rather than just relying on the oldest fossils of clades. In addition, both approaches assume that fossil records of different clades in the phylogeny are all the product of the same underlying fossil sampling rate, even though this rate has been shown to differ strongly between higher-level taxa. We here develop a flexible new approach to Bayesian node dating that combines advantages of traditional node dating and the fossilized birth-death model. In our new approach, calibration densities are defined on the basis of first fossil occurrences and sampling rate estimates that can be specified separately for all clades. We verify our approach with a large number of simulated datasets, and compare its performance to that of the fossilized birth-death model. We find that our approach produces reliable age estimates that are robust to model violation, on par with the fossilized birth-death model. By applying our approach to a large dataset including sequence data from over 1000 species of teleost fishes as well as 147 carefully selected fossil constraints, we recover a timeline of teleost diversification that is incompatible with previously assumed vicariant divergences of freshwater fishes. Our results instead provide strong evidence for trans-oceanic dispersal of cichlids and other groups of teleost fishes. (Keywords: node dating; calibration density; relaxed molecular clock; fossil record; Cichlidae; marine dispersal)

---

In phylogenetic analyses, molecular sequence data are commonly used to infer not only the relationships between species, but also the divergence times between them. The estimation of divergence times in a phyogenetic context is usually based on an assumed correlation between the age of species separation and the number of observed genetic differences, i.e. a “molecular clock” (Zuckerkandl and Pauling 1962). Evidence for the existence of molecular clocks initially derived from relative rate tests (Sarich and Wilson 1967) and has since been corroborated by a large body of literature (e.g. Wilson et al. 1977; Bromham and Penny 2003). However, it has been shown that the rate of the molecular clock often differs between lineages (Drummond et al. 2006) and that it can depend on factors including body size, metabolic rate, and generation time (Martin and Palumbi 1993; Nabholz et al. 2008).

To allow the estimation of absolute divergence dates from sequence data, a calibration of the rate of the molecular clock is required. This calibration can be obtained from serially sampled DNA sequences, if the range of sampling times is wide enough to allow accumulation of substantial genetic differences between the first and last sampling event (Drummond et al. 2003), which is often the case for rapidly evolving viruses (Faria et al. 2014; Smith et al. 2009; Gire et al. 2014). However, for macroevolutionary studies aiming to estimate divergence times of eukaryotic organisms on the order of tens or hundreds of million years, genetic differences accumulated between sampling times are negligible and other sources of calibration information are required. Commonly, the age of the oldest known fossil of a given clade is then used to calibrate the age of this clade, an approach often referred to as “node dating” (Ronquist et al. 2012; Grimm et al. 2015). However, due to the incompleteness of the fossil record, clade origin will almost always predate the preservation of its oldest known fossil. As a result, fossils can provide absolute minimum clade ages, but are usually less informative regarding the maximum ages of clades (Benton and Donoghue 2007). In a Bayesian framework for phylogenetic time-calibration, the uncertainty regarding clade ages can be accomodated by the specification of “calibration densities” (also referred to as “node age priors”) with a hard lower bound set to the age of the earliest fossil record and a soft upper bound as provided by exponential, lognormal, or gamma distributions. Unfortunately, the optimal parameterization of these distributions is usually unknown but has been shown to have a strong influence on the resulting age estimates (Ho and Phillips 2009). In addition, the effect of inaccurate calibration densities can only partially be corrected with larger molecular datasets (Yang and Rannala 2006).

Other shortcomings of node dating have been identified. As described by Heled and Drummond (2012), calibration densities interact with each other and with the tree prior to produce marginal prior distributions of node ages that may differ substantially and in unpredictable ways from the specified calibration density. The application of recently-introduced calibrated tree priors can compensate for this effect, but becomes computationally expensive when more than a handful of calibrations are used in the analysis (Heled and Drummond 2015). Node dating has also been criticized for ignoring most of the information from the fossil record, as only the oldest known fossils of each clade are used to define calibration densities (Ronquist et al. 2012, but see Marshall 2008; Claramunt and Cracraft 2015). Furthermore, node dating relies on the correct taxonomic assignment of fossils to clades, and may produce misleading age estimates when fossils are misplaced on the phylogeny (Marshall 2008; Ho and Phillips 2009; Forest 2009).

As alternatives to node dating, two approaches have recently been developed. In “total-evidence” dating, fossils are not explicitely assigned to any clades, but are instead included as terminal taxa. The position of these tips is determined as part of the phylogenetic analysis, based on morphological character data that are required for all included fossils and at least some of the extant taxa (Pyron 2011; Ronquist et al. 2012). Branch lengths, and thus divergence times between extant and extinct species are inferred under the assumption of a “morphological clock”, usually based on the Mk model of Lewis (2001) (Pyron 2011; Beck and Lee 2014; Arcila et al. 2015). The total-evidence approach is conceptually appealing as it is able to account for uncertainty in the phylogenetic position of fossils, and allows a more complete representation of the fossil record than node dating. However, this approach has been found to result in particularly ancient age estimates and long “ghost lineages” when applied to empirical data sets, leading some authors to question the suitability of morphological clocks for phylogenetic time-calibration (Beck and Lee 2014; Arcila et al. 2015; O’Reilly et al. 2015). The developments of more realistic sampling schemes (Höhna et al. 2011) and advanced models of morphological character evolution (Wright et al. 2015) are likely to improve age estimates obtained with total-evidence dating, but have so far been applied only rarely (Klopfstein et al. 2015; Zhang et al. 2015). Importantly, due to the requirement of a morphological character matrix, total-evidence dating is limited to groups that share sufficient numbers of homologous characters (Grimm et al. 2015) so that its application is practically not feasible for higher-level phylogenies combining very disparate taxa from different taxonomic orders or classes.

The “fossilized birth-death (FBD) process” (Heath et al. 2014) provides an elegant framework in which fossils are used as terminal taxa and thus more than the oldest fossil can be used for each clade. In this model, fossils as well as extant taxa are considered as the outcome of a common process based on the four parameters speciation rate λ, extinction rate *μ*, proportion of sampled extant taxa *ρ*, and the fossil “sampling rate” *ψ* It is assumed that the fossils that are ultimately sampled and included in the study have been preserved along branches of the complete species tree following a constant-rate Poisson process. Unlike in total-evidence dating, a morphological character matrix is not required to place fossil taxa in the phylogeny (but can be used for this purpose; Gavryushkina et al. 2015; Zhang et al. 2015). Instead, fossils are assigned to clades through the specification of topological constraints. The FBD process was first implemented in the Bayesian divergence time estimation program FDPPDIV (Heath et al. 2014), which requires the specification of a fixed tree topology, point estimates of fossil ages, and constant rates of diversification and sampling throughout the tree. All these limitations have subsequently been overcome in the implementations of the FBD process as a tree prior in the “Sampled Ancestors” package for BEAST (Gavryushkina et al. 2014; Bouckaert et al. 2014) and in the software MrBayes (Huelsenbeck and Ronquist 2001; Ronquist and Huelsenbeck 2003; Zhang et al. 2015). These implementations allow to specify priors on fossil ages to account for the often large uncertainties associated with them as well as the specification of time intervals within which rates are assumed constant, but between which they are free to vary. However, a limitation that remains also in newer FBD implementations is the assumption that all clades existing in a given time interval are subject to the same rates of diversification and fossil sampling. Especially in higher-level phylogenies, this assumption is unlikely to be met, as substantial clade-specific differences in these rates have been identified in many groups (Foote and Sepkoski 1999; Alfaro et al. 2009; Jetz et al. 2012; also see Supplementary Table S1), suggesting that time estimates obtained on the basis of this assumption may be misleading. The FBD model further assumes that the fossils included in the analysis represent either the complete set or a random sample of the known fossil record of a clade. However, the use of a complete or randomly sampled representation of the fossil record may be impractical with higher-level phylogenies due to the enormous number of fossil occurrences known for many higher taxa, e.g. for mammals (> 90 000), birds (> 5 000), and insects (> 40 000; www.paleobiodb.org). Presumably as a consequence of these difficulties, node dating has remained popular despite its drawbacks, and was applied in all recent phylogenomic time-tree analyses of groups above the order level (mammals: dos Reis et al. 2014; birds: Jarvis et al. 2014; Prum et al. 2015; insects: Misof et al. 2014).

Here, we develop a new approach for Bayesian phylogenetic divergence-time estimation that is based on node dating, but infers the optimal shape of calibration densities from a combination of the first fossil occurrence age of a given clade and independently assessed estimates of the sampling rate and the diversification rates. This approach therefore overcomes a major problem of node dating, the fact that calibration densities are often chosen arbitrarily despite their strong influence on the resulting age estimates (Heath et al. 2014). Our approach is suitable for divergence-time estimation in higher-level phylogenies combining groups with different sampling or diversification characteristics, as parameters can be specified independently for each clade. We have implemented our method in a new package for BEAST called “CladeAge”, and we will refer to calibration densities obtained with it as “CladeAge calibration densities” throughout the manuscript. Using a wide range of simulations, we assess the optimal scheme by which to select clades for calibration, and we show that the application of CladeAge calibration densities can result in age estimates comparable or better than those produced with the FBD model if the input rate estimates are correctly specified and only the oldest fossil of each clade is used for calibration.

We use our new approach together with a large and partially newly-generated molecular data set of 1187 teleost fishes to address the long-standing question whether freshwater cichlid fishes from India, Madagascar, Africa and the Neotropics diverged before or after continental separation (Chakrabarty 2004; Sparks and Smith 2005; Genner et al. 2007; Azuma et al. 2008; Friedman et al. 2013; McMahan et al. 2013). By rigorous examination of the paleontological literature, we identify as many as 147 fossil calibrations that can be reliably assigned to monophyletic clades in our phylogeny. To our knowledge this is the largest number of fossil calibrations thus far applied in a single time-tree analysis, and it suggests that studies using phylogenetic divergence-time estimation commonly underutilize the fossil record. Our results strongly support divergence of freshwater fishes long after continental separation, implying multiple marine dispersal events not only in cichlid fishes but also in other freshwater groups included as outgroups in our phylogeny.

## CladeAge calibration densities

### Calculating CladeAge calibration densities

Here, our goal is to design calibration densities that reproduce the probability density for a clade originating at time *t_o_,* given the age of its oldest fossil *t_f_*. To estimate this probability density, we assume that the probability density of a clade being *t* time units older than it oldest fossil is identical to the probability density *f_s_(t)* of the oldest fossil being *t* time units younger than the clade origin. This is equivalent to assuming a uniform prior probability distribution for the age of the clade, which is justified for calibration densities, as these probability densities will be multiplied with a (non-uniform) tree prior at a later stage, during the divergence time analysis. Thus, any non-uniform prior assumptions about the clade origin can be incorporated via the tree prior. We further assume that speciation, extinction, and fossil sampling are all homogeneous Poisson processes with rates *⋋, μ*, and *ψ*, respectively.

For a single lineage that does not speciate or go extinct, the probability to remain unsampled until time *t_1_* is *p_u_(t_1_) = e^−ψt_1_^*, while the probability of being sampled at least once during the same period is *p_s_(t_1_) = 1* — *e^−ψt_1_^*. Thus the probability of not being sampled before time *t*_1_, but then being sampled before time *t_2_ > t_1_* is

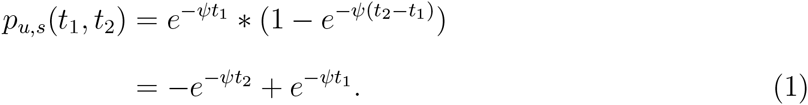

The probability density for the clade being sampled for the first time exactly at time *t_1_* is then

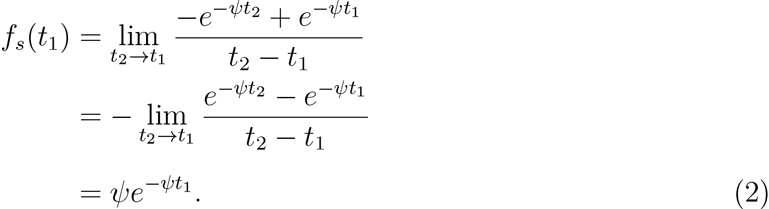

If we now allow for the possibility that the lineage has diversified into *N* species extant at time *t_1_*, then the probability of the clade not being sampled before time *t_1_* is *p_u_(t_1_) = ^e-ψS(t_1_)^*, where *S(t_1_)* is the sum of lineage durations between clade origin and time *t_1_* (Foote et al. 1999). The probability of no lineage being sampled before time t_1_, but at least one lineage being sampled before time *t*2 is then

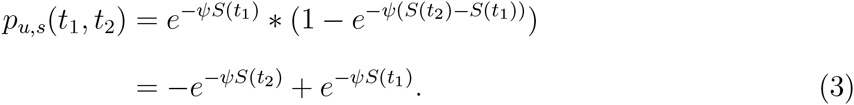

In this case, the probability density for the clade being sampled for the first time exactly at *t_1_* is

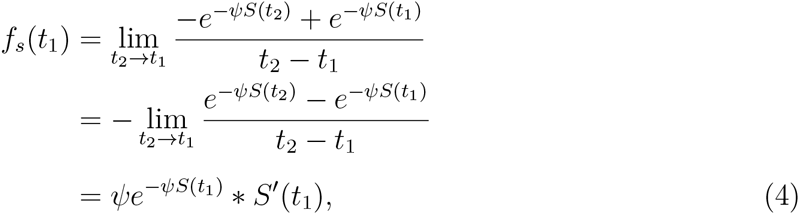

where the first derivative of *S(t)* at time *t_1_* is

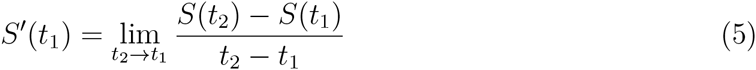

By ignoring the possibility of speciation or extinction between *t_1_* and *t_2_* (which is justified at the limit *t_2_ → t_1_*), we get *S(t_2_) = S(t_1_) + N * (t_2_ —1_1_)* and thus *S’(t_1_) = N*, which gives us

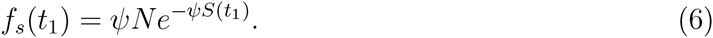

If we now take into account the stochastic nature of *S* as a variable resulting from a birth-death process with parameters *⋋* and *μ*, we have to rewrite Equation 6 as

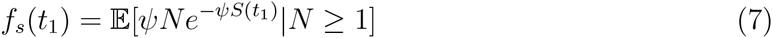

where we condition on the survival of at least one species at time *t_1_*, which is necessary to allow sampling at this time.

Unfortunately, we can not solve *f_s_(t_1_)* analytically. To approximate *f_s_(t_1_),* CladeAge generates 10 000 birth-death trees based on estimates of the speciation rate *⋋* and the extinction rate *μ*, infers *S(t_1_)* in each of these trees as the sum of all branch lengths between clade origin and *t_1_*, and calculates *ψNe^−ψS(t_1_)^* if the birth-death process resulted in *N ≥ 1*. According to the law of large numbers, the mean of a large sample converges to its expected value, therefore the probability density *f_s_(t_1_)* can be approximated by the mean of all values calculated for *ψNe^−ψS(t_1_)^*. This process is repeated for 100 time points evenly spaced between 0 and a maximum time *t_max_,* which is predetermined so that *p_u_(t_max_)* ≈ *0.001 * p_u_(0).* The probability density *f_s_(t)* for times *t* in between two of the 100 time points is estimated through interpolation from the probability densities of the two neighbouring time points, using linear regression. For all times larger than *t_max_,* probability densities are approximated by a scaled exponential distribution that is calculated on the basis of the two largest time points and their respective probability densities *f_s_(t).* Finally, all estimates of probability densities are scaled so that the total probability mass becomes 1.

The calculation of calibration densities, as described above, requires estimates of the fossil sampling rate, as well as of the speciation and extinction rates, which can be obtained externally, from the fossil record alone (Silvestro et al. 2014; Starrfelt and Liow 2015), or from a combination of fossil and phylogenetic information (Alfaro et al. 2009; Stadler 2011; Rabosky 2014). As diversification is commonly parameterized as “net diversification” (*⋋ — μ)* and “turnover” (*μ/⋋*), and researchers often have greater confidence in estimates of net diversification and turnover than in those for speciation and extinction rates (Beaulieu and Donoghue 2013), our method accepts input in these units, and calculates *λ* and *μ* from it. Uncertainty in the three parameters net diversification, turnover, and sampling rate can be expressed by specifying minimum and maximum values and is accounted for by randomly drawing from the specified ranges, for each of the 10 000 birth-death trees generated to estimate estimated probability density *f_s_*. Examples demonstrating the shape of CladeAge calibration densities, based on exactly known (A) or uncertain ages of the first fossil record (B), are shown in Figure 1.

**Figure 1:**
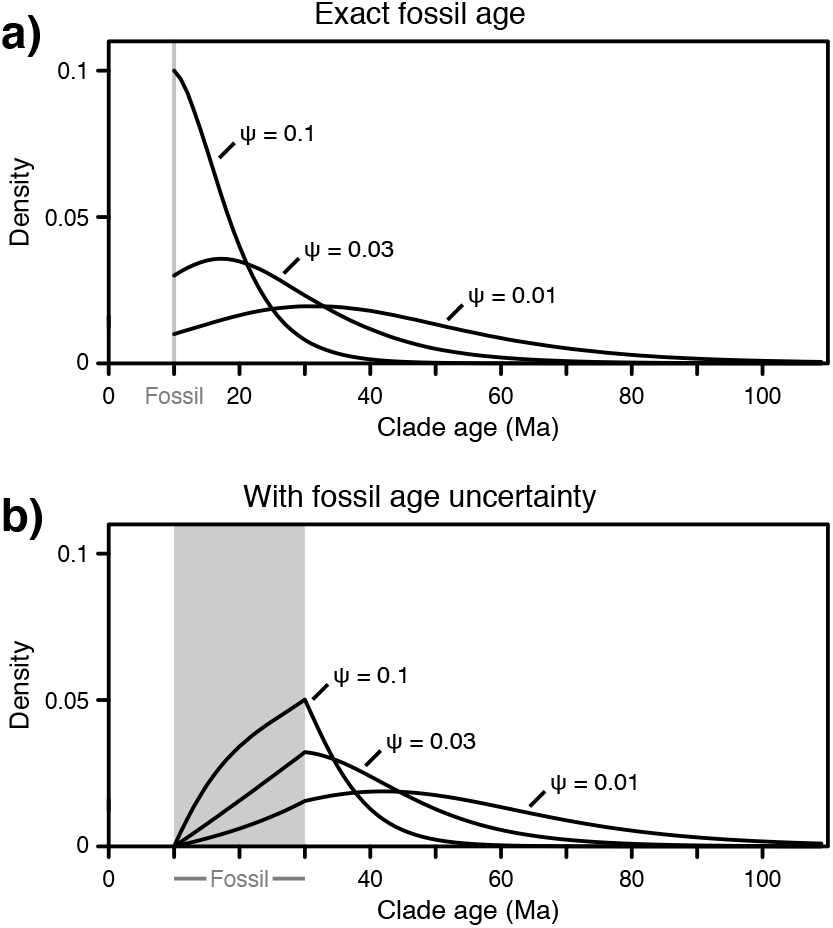
Exemplary CladeAge calibration densities. Probability densities for the age of a clade for which the earliest fossil is known to be exactly 10 myr old (a), or assumed to be between 10 and 30 myr old, with a uniform fossil age probability within this range (b). The gray area in b) indicates the fossil age uncertainty. Speciation rate and extinction rates are assumed to be *λ* = 0.08 and *μ* = 0.04, and sampling rates *ψ* are as indicated.

### Calibration schemes for the use of CladeAge calibration densities in phylogenetic divergence-time estimation

Under the assumption of constant rates of diversification and sampling as well as a uniform prior probability for node ages, CladeAge calibration densities approximate the probability density for the age of a clade, given the age of the oldest fossil record of this clade. These probability distributions are therefore suitable as constraints on clade ages in Bayesian divergence-time estimation. However, in practice, it may not always be clear which clades should be used for time calibration: All clades with fossil records? Only those clades with stem-group fossils? Or only those with a crown-group fossil record? If a fossil represents the earliest record of not only one clade, but of multiple nested clades, CladeAge calibration densities could be used as time constraints for all these clades (we refer to this as “scheme A”), only for the most inclusive of these clades (“scheme B”), or only for the least inclusive clade (“scheme C”). As scheme B would allow one or more of the clades to appear younger than the fossil itself, it seems reasonable to specify, in addition to the CladeAge calibration density for the most inclusive clade, the fossil age as a strict minimum age for the least inclusive clade when using this scheme. Furthermore, if two sister clades both possess a fossil record, they could both be used to constrain the age of the two clades. However, as the ages of the two clades are necessarily linked by their simultaneous divergence, two time constraints would be placed on one and the same node. Instead, it may seem more intuitive to use only the older of the two fossils for time-calibration and disregard the younger fossil (“scheme D”). However, in contrast to traditional node dating, where maximally one calibration density is placed on each node, the model used to calculate CladeAge calibration densities considers each clade individually, and could thus be biased if the selection of clades for calibration is based on information about their sister clade. Figure 2a illustrates the four different calibration schemes.

**Figure 2:**
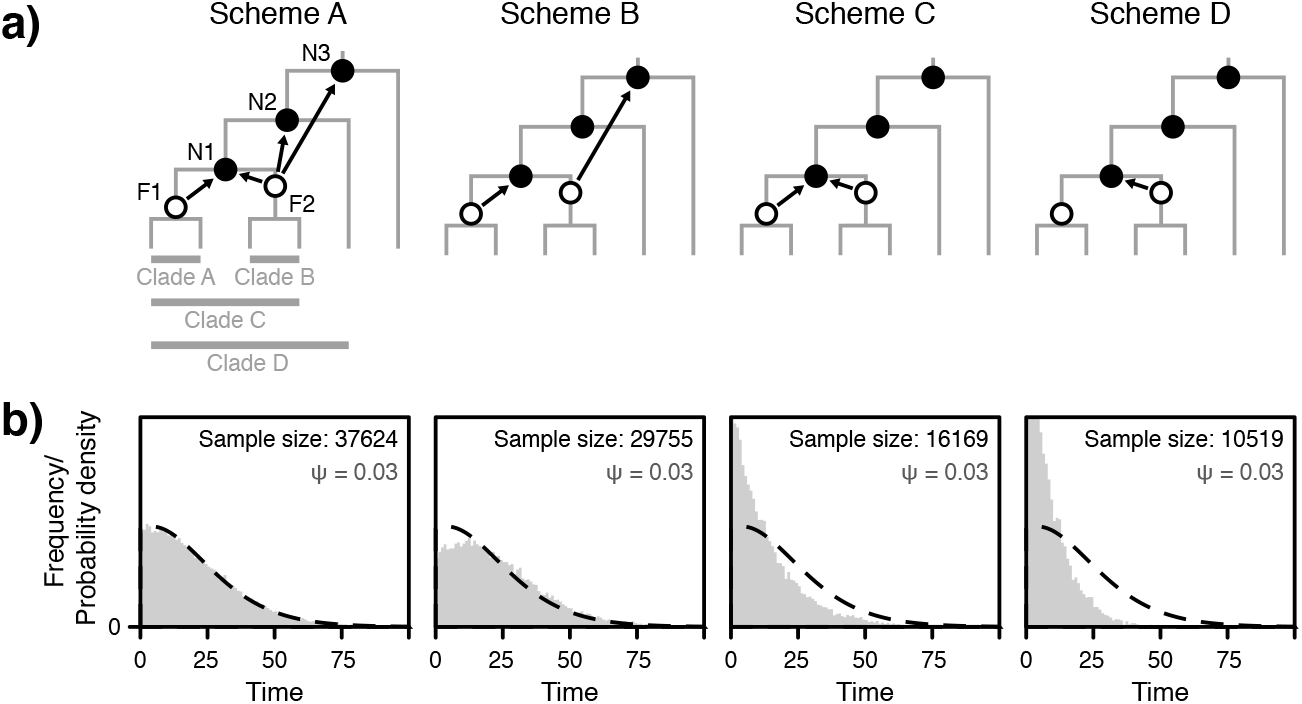
Four alternative calibration schemes for CladeAge calibration densities. a) Assume that fossils F1 and F2 (represented by white circles) can be assigned to the stems of clade A and clade B, respectively, based on their morphology. As fossil F2 represents the first fossil record not only of clade B, but also of the more inclusive clades C and D, it could be used to constrain the stem age of these three clades, and thus nodes N1, N2, and N3. In addition, since F1 can be used to constrain the stem age of clade A, and F2 can be used to constrain the stem age of clade B, to different constraints could be placed on the divergence of clade A and B, represented by node N1. In scheme A, both fossils are used to constrain the stem ages of all clades, for which these fossils represent the earliest record. In schemes B and C, each fossil constrains only the most inclusive clade (scheme B), or only the least inclusive clade (scheme C), for which it represents the first occurrence. In scheme D, each fossil constrains just one clade that is drawn at random if multiple options exist. Scheme E is similar to scheme A except that only the older one of two fossils in two sister clades is used to constrain the common divergence event. b) Comparison of waiting times between clade origin and first fossil occurrence. Waiting times between clade origin and first fossil occurrence were recorded from 10 000 simulated phylogenies with three different sampling rates (*ψ* = 0.1, *ψ* = 0.03, *ψ* = 0.01), using schemes A-E, and a clade age threshold of 0.9. The frequency distributions of binned waiting times are shown in gray, and CladeAge probability density distributions for the same settings are indicated with dashed black lines. The total number of waiting times sampled is given in each plot.

As CladeAge calibration densities approximate the probability densities of clade ages conditional on the age of the first fossil record of this clade, they are also expected to approximate frequency distributions of observed waiting times between the origin of a clade and the appearance of the first fossil record of this clade in a sufficiently large sample of simulated phylogenetic trees. Since these waiting times can be sampled according to the above four schemes, we can determine the optimal calibration scheme by comparison of waiting time frequency distributions with CladeAge calibration densities. We simulated three times 10 000 pure-birth phylogenies with a speciation rate *⋋ = 0.04* and a root age *t_root_* randomly drawn from a uniform distribution between 20 and 200 time units, conditioned on the survival of exactly 100 extant species. Assuming a Poisson process of fossil sampling, we added simulated fossil records to the branches of each of these trees, with three different sampling rates *ψ = 0.1, 0.03, 0.01.* Applying the above four calibration schemes (A-D) independently, we recorded waiting times between a clade’s origin and the age of its oldest fossil in each simulated phylogeny.

Waiting time frequency distributions recorded from relatively young clades can be biased by the fact that only those waiting times shorter than the clade age can be recorded (otherwise the clade did not preserve at all). To assess the degree of this effect, we repeated this analysis, counting only waiting times for clades with a time of clade origin *t_o_* above one out of four thresholds: *t_o_ ≥ 0* (all clades included), *t_o_ ≥ 0.5) × t_root_, t_o_ ≥ 0.9 × t_root_,* and *t_o_ = 1 × t_root_* (including only the two clades descending from the root, per simulated phylogeny). With the strictest clade age threshold of *t_o_ = 1 × t_root_,* the same two waiting times per phylogeny are recorded with schemes A and B if both clades descending from the root have produced fossils. This is because the root node represents the oldest node that can be constrained with fossils in these clades, and thus waiting times between the root and these fossils are recorded with both schemes A and B. If further divergence events occurred between the root and the fossil, the root does not represent the youngest node that can be constrained with the fossil, and thus, the waiting time between the root and the fossil are not recorded with schemes C and D (see Fig. 2a). Differences between schemes A and B become apparent with less strict clade age thresholds, when also clades are included that do not represent the oldest possible clade to be constrained with a given fossil.

Figure 2b shows comparisons between waiting time frequency distributions and CladeAge calibration densities for a clade age threshold of *t_o_ ≥ 0. 9 × t_root_,* which is sufficiently young to show differences between all schemes, but still old enough to be affected only minimally by the bias described above. Comparisons for all other tested clade age thresholds are shown in Supplementary Figure S1. Taken together, these results show that waiting time frequency distributions deviate from the respective CladeAge distribution in most comparisons, and the degree of disagreement depends on sampling rate 0, on the clade age threshold, and on the applied scheme (A-D). However, for all but the youngest clade age thresholds, scheme A produces a frequency distribution that is virtually identical in shape to the distribution of CladeAge calibration densities. This suggests that when CladeAge calibration densities are used for time calibration, they should strictly be applied to constrain all clades for which a given fossil represents the first occurrence, even if the same fossil is used to constrain multiple nodes, and even if more than one constraint is placed on one and the same node.

## Testing CladeAge Calibration Densities with Simulated Phylogenies

To more extensively compare the performance of the four different calibration schemes A to D, we simulated phylogenetic data sets including fossil records and sequence alignments, and used CladeAge calibration densities to estimate clade ages in BEAST v.2.1.3. For comparison, we also used the same generated data sets to estimate clade ages with the FBD process implemented in the Sampled Ancestors (Gavryushkina et al. 2014) package for BEAST.

### Generation of data sets

Phylogenetic data sets of trees and fossil records were generated as decribed above with sampling rates *ψ = 0.1, 0.03, 0.01,* a root age between 20 and 200 time units, and a net diversification *⋋ — μ* of 0.04, however, species turnover was now modeled with rate *μ/⋋ = 0.5* (thus using *⋋ = 0.08* and *μ = 0.04).* If the time units used in these simulations are considered to be million years, the sampling and diversification rates used here are comparable to those found in empirical data sets (Jetz et al. 2012; Stadler and Bokma 2013; Rabosky et al. 2013; Supplementary Table S1). In separate sets of simulations, branch-specific substitution rates were modeled either with an uncorrelated molecular clock (Drummond et al. 2006), or with an autocorrelated molecular clock that accounts for the heritability of factors influencing rate variation (such as body mass, longevity, and generation time; Nabholz et al. 2008; Amster and Sella 2016) and may therefore model rate evolution more realistically than the uncorrelated molecular clock (Lepage et al. 2007). For both types of branch rate variation, we used a mean rate of *4 × 10^−3^* substitutions per site per time unit and a variance parameter of *1.6 ×* 10^−5^. Branch-rate autocorrelation was simulated with the Cox-Ingersoll-Ross (CIR) process of Lepage et al. (2006), using a decorrelation time of 100 time units. The branch lengths and substitution rates were used to simulate sequence evolution of 3000 nucleotides according to the unrestricted empirical codon model of Kosiol et al. (2007). For each of two clock models and each of the three sampling rates, we generated 50 replicate data sets. An example of a data set simulated with these settings is illustrated in Supplementary Figure S2.

### Phylogenetic divergence-time estimation

For each of the replicate data sets, the simulated phylogenetic trees were reconstructed, and for each clade in each reconstructed phylogeny, the oldest fossil occurrence was identified. CladeAge calibration densities were calculated for these fossils based on the parameters used in simulations (λ = 0.08, *μ* = 0.04, and *ψ* = 0.1, 0.03, 0.01), and used to constrain node ages according to calibration schemes A to D in divergence-time estimation with BEAST. All sequence alignments were divided into three partitions according to codon position, and for each partition, we used the reversible-jump based substitution model of Bouckaert et al. (2013) with four gamma-distributed rate categories. For all simulated data sets, we used the lognormal relaxed molecular clock (Drummond et al. 2006) for divergence-time estimation. To account for extinction in the diversification process, we used the birth-death tree prior of Gernhard (2008) with uninformative prior distributions for the birth rate and the relative death rate. We used the reconstructed simulated tree as a starting tree in all analyses, and fixed the tree topology by disallowing all topological changes. For each analysis, 50 million Monte Carlo Markov Chain (MCMC) steps were carried out, which was always sufficient for convergence.

For the analysis of the same data sets with the FBD model implemented in the Sampled Ancestors package for BEAST, we used settings as described above, except that between 100 and 400 million MCMC steps were required for convergence. For comparability with age estimates based on CladeAge calibration densities, we used only the oldest fossils for each clade, and we fixed the values of diversification rates to those used to generate the data set. In separate sets of analyses, however, we fixed the sampling proportion according to the sampling rate used in simulations, or allowed the sampling proportion to be estimated. We also fixed the tree topology of all extant species, while at the same time allowing fossil taxa to attach anywhere whithin the clade (including its stem) to which they were assigned. This was done by using instances of “CladeConstraint”, a new type of topological constraint for BEAST introduced as part of the Sampled Ancestors package (Gavryushkina et al. 2014), with which ingroups and outgroups can be defined for a given clade, and taxa not listed in either of these groups are free to appear in either of them. For each clade, we specified CladeConstraints that place all extant taxa and fossils of this clade within the ingroup and all other extant taxa in the outgroup, thus allowing fossils from parent clades to appear outside or within this clade. As the starting tree, we used the reconstructed simulated tree but reattached each clade’s oldest fossil (provided that it had any) to its stem with an additional branch. All BEAST input files used for the analysis of simulated datasets are provided as Supplementary Data S1.

### Results with simulated phylogenies

Our simulations produced phylogenetic trees with root heights between 52.2 and 163.8 time units, with a median height of 84.8 time units. Mean branch rates per tree were between 2.2 × 10^−3,^ and 6.6 × 10^−3,^ (median 3.9 × 10^−3^) substitutions per time unit with branch rate variances between 7.8 × 10^−7^ and 4.4 × 10^−5^ (median 7.1 × 10^−6^), resulting in 5280 to 15090 (median 8289) nucleotide substitutions. The sequence alignments contained between 2217 and 2836 (median 2568) variable sites and between 1724 and 2646 (median 2231) parsimony-informative sites, out of a total of 3000 sites per alignment. Simulated fossil records consisted of 165 to 380 (median 240.5) fossils when generated with a sampling rate of *ψ* = 0. 1, 40 to 123 (median 74.5) fossils with *ψ* = 0.03, and 10 and 49 (median 24) fossils when a sampling rate of *ψ* = 0.01 was applied (see Supplementary Figure S2 for an illustration). Discarding fossils that did not represent the oldest fossil of any clade left between 75 and 113 (median 94) fossils when the sampling rate was *ψ* = 0.1, 30 to 71 (median 47) fossils with 0 = 0.03, and 9 to 33 (median 19.5) fossils with *ψ* = 0.01. Figure 3a shows the mean number of fossil constraints in 50 simulated data sets, per bin of 20 time units. The number of fossils available as time constraints decreases with bin age, a direct result of the fact that younger time bins contain an overall larger sum of lineage durations.

**Figure 3:**
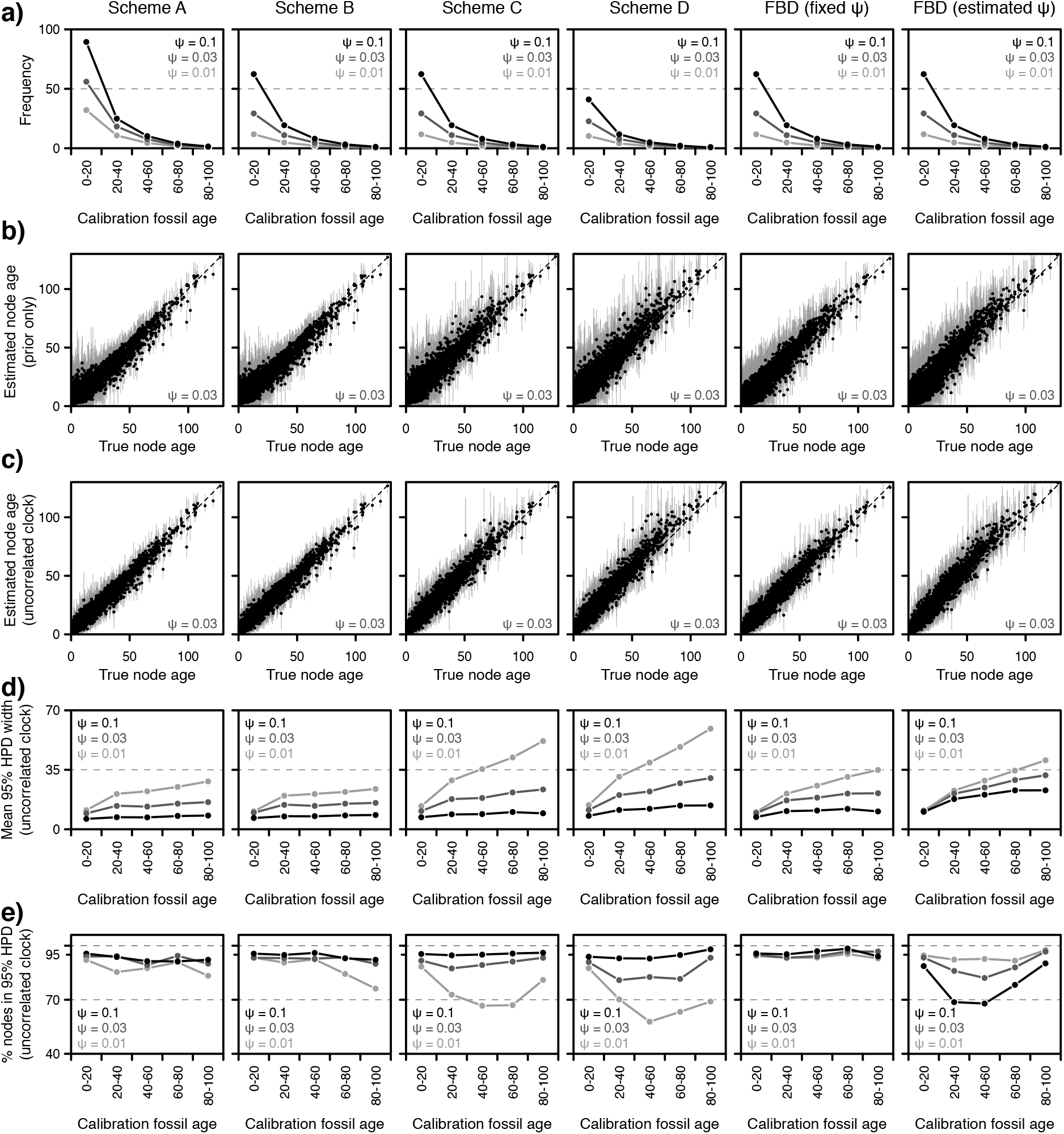
Phylogenetic divergence-time estimation with simulated data sets, using four CladeAge calibration schemes and the FBD process. Results are based on 50 simulated phylogenetic trees and sequence data, and fossil records simulated with three different sampling rates for each phylogeny. a) The mean number of fossil constraints used with each scheme, sorted into bins of 20 time units according to fossil age. For schemes B, C, and the FBD model, this number is identical to the number of fossils. In scheme A, some fossils are used for multiple constraints, and in scheme D, not all fossils are used (see Fig. 2). b) Estimated node ages with MCMC sampling from the prior alone, when the fossil record was simulated with the intermediate sampling rate 0 = 0.03. Node age comparisons based on other sampling rates (*ψ* = 0.1 or *ψ* = 0.01) are shown in Supplementary Figure S3. c) As b, but using MCMC sampling from the posterior, with sequence data generated with uncorrelated branch rates. Results for data sets with autocorrelated branch rates are shown in Figure S3. d) Mean width of 95% HPD intervals, when using MCMC sampling from the posterior with datasets generated with uncorrelated branch rates. Results are given in bins of 20 time units according to the true node age. e) Percentage of age estimates for which the 95% HPD interval includes the true node age, when sampling from the posterior and using datasets generated with uncorrelated branch rates. As in d), results are presented in bins of 20 time units according to the true node age. See Supplementary Tables S10-S12 for summary statistics for the full set of analyses.

Comparisons of estimated and true node ages are shown in Figure 3b-c, for all analyses of data sets generated with the intermediate sampling rate of *ψ* = 0.03 and the uncorrelated clock model (see Supplementary Figure S3 for results obtained with *ψ* = 0.1 or *ψ* = 0.01, or with the autocorrelated clock model). The difference between results based on MCMC sampling from the prior only (Fig. 3b) and results based on the posterior (Fig. 3c) is most pronounced for young clades where 95% highest posterior density (HPD) intervals (indicated with gray bars in Fig. 3b-c) are much wider when the MCMC sampled from the prior only. This suggests that in combination with a relaxed clock model, sequence data is most informative to determine the age of young nodes, but that the age estimates of older nodes are primarily determined by the specified prior probabilities.

Following Heath et al. (2014) and Gavryushkina et al. (2014), we describe the age estimates for simulated phylogenies with two summary statistics, the mean width of 95% HPD intervals and the percentage of 95% HPD intervals that include the true node age. Shorter 95% HPD intervals indicate greater precision, and the percentage of 95% HPD intervals that include the true node age serves to assess the accuracy of age estimates. If the model used to generate the data is identical to that assumed for divergence-time estimation, and if MCMC sampling has completely converged, 95% of the 95% HPD intervals are expected to include the true node age. For CladeAge analyses of data sets generated with uncorrelated branch rates, a nearly identical model was used for simulation and inference, and the resulting percentage of 95% HPD intervals containing the true node age can therefore serve as an indicator of the optimal calibration scheme to be used with CladeAge. In contrast, the model used in analyses with the FBD differs to a greater extent from the model used to generate data sets, as the FBD model assumes that all, or a randomly sampled set of fossils of a clade are used for calibration, whereas our data sets were reduced to contain only the oldest fossils of each clade. Thus, for FBD analyses of data sets generated with uncorrelated branch rates, the two summary statistics allow to assess the robustness of the FBD model to a violation of the assumed fossil record representation. In addition, results for data sets generated with autocorrelated branch rates indicate the degree of robustness of both CladeAge and FBD analyses to violations of the clock model assumed for divergence-time estimation.

For all analyses with CladeAge as well as FBD analyses in which the sampling rate was allowed to be estimated, the two summary statistics are listed in Table 1. For data sets generated with the uncorrelated clock model, these results are also illustrated in bins of 20 time units in Figure 3d-e (detailed results for all analyses are given in Supplementary Tables S10-S12).

**Table 1.** Estimated node ages for simulated phylogenies, based on four CladeAge calibration schemes and the FBD process. Divergence-time estimation was based on MCMC sampling from prior probabilities alone, or in combination with the likelihood of sequence data simulated with uncorrelated or autocorrelated branch rates. Fossil records were simulated with three different sampling rates *ψ* = 0.1, 0.03, 0.01. For the FBD process, results are shown for analyses in which the sampling rate *ψ* was either fixed or allowed to be estimated.

| **Mean 95% HPD width:** |  |  |  |  |  |  |  |
| --- | --- | --- | --- | --- | --- | --- | --- |
| Clock model | $\psi$ | Scheme A | Scheme B | Scheme C | Scheme D | FBD (fixed $\psi$) | FBD (est. $\psi$) |
| prior only | 0.1 | 9.34 | 10.30 | 11.55 | 14.07 | 11.84 | 20.01 |
| prior only | 0.03 | 17.42 | 18.45 | 21.84 | 24.27 | 18.11 | 22.30 |
| prior only | 0.01 | 25.28 | 24.13 | 33.26 | 35.72 | 22.03 | 23.66 |
| uncorrelated | 0.1 | 6.41 | 6.91 | 7.55 | 9.00 | 8.20 | 12.68 |
| uncorrelated | 0.03 | 10.63 | 11.06 | 12.91 | 14.30 | 11.90 | 14.39 |
| uncorrelated | 0.01 | 14.08 | 13.19 | 18.67 | 19.99 | 13.71 | 14.77 |
| autocorrelated | 0.1 | 4.56 | 4.82 | 5.10 | 6.03 | 5.75 | 8.66 |
| autocorrelated | 0.03 | 6.77 | 6.92 | 7.83 | 8.73 | 7.84 | 9.44 |
| autocorrelated | 0.01 | 8.49 | 8.00 | 11.39 | 12.63 | 8.76 | 9.61 |

| **Percentage of 95% HPD intervals containing the true node age:** |  |  |  |  |  |  |  |
| --- | --- | --- | --- | --- | --- | --- | --- |
| Clock model | $\psi$ | Scheme A | Scheme B | Scheme C | Scheme D | FBD (fixed $\psi$) | FBD (est. $\psi$) |
| prior only | 0.1 | 95.2 | 96.7 | 96.1 | 95.6 | 98.2 | 93.0 |
| prior only | 0.03 | 94.8 | 94.4 | 93.2 | 91.5 | 97.0 | 96.1 |
| prior only | 0.01 | 92.6 | 94.3 | 89.5 | 88.5 | 96.1 | 96.5 |
| uncorrelated | 0.1 | 94.9 | 95.4 | 95.3 | 93.8 | 95.8 | 83.7 |
| uncorrelated | 0.03 | 93.7 | 93.2 | 90.9 | 88.3 | 94.7 | 91.1 |
| uncorrelated | 0.01 | 90.5 | 92.4 | 83.9 | 82.0 | 94.1 | 94.0 |
| autocorrelated | 0.1 | 87.9 | 87.3 | 86.4 | 81.9 | 84.4 | 63.5 |
| autocorrelated | 0.03 | 76.8 | 75.7 | 69.3 | 62.7 | 72.2 | 63.8 |
| autocorrelated | 0.01 | 66.2 | 70.3 | 57.0 | 54.3 | 70.3 | 67.7 |

Among the four calibration schemes A to D, scheme A produced the shortest 95% HPD intervals with data sets based on *ψ* = 0. 1 or *ψ* = 0.03, regardless of whether the MCMC was set to sample from the prior only, or from the posterior, and both with data sets generated with uncorrelated or autocorrelated branch rates. At the same time, the percentage of true node ages included in 95% HPD intervals obtained with scheme A is closer to the expected value of 95% than that of any other calibration scheme. In contrast, scheme B performed slightly better than scheme A for data sets with the lowest sampling rate *ψ* = 0.01, as indicated by shorter 95% HPD intervals and a greater percentage of true node ages included within them. However, when scheme B was used for the analysis of data sets generated with uncorrelated branch rates, the accuracy of age estimates decreased with node age, and for nodes with a true age between 80 and 100 time units, only 76.2% of the 95% HPD intervals contained their true age (Fig. 3e, Supplementary Table S11).

In all cases, MCMC sampling from the posterior decreased the mean width of 95% HPD intervals, compared to analyses using the prior probability alone. The percentage of 95% HPD intervals containining the true node age remained comparable between analyses based on the prior probability alone (92.6-95.2% with scheme A) and analyses using the posterior (90.5-94.9% with scheme A) for data sets generated with uncorrelated branch rates. However, for data sets generated with autocorrelated branch rates, the percentage of 95% HPD intervals containining the true node age decreased substantially when the posterior was used for MCMC sampling (66.2-87.9% with scheme A; Table 1).

Overall, FBD analyses with a fixed sampling rate produced very similar summary statistics to CladeAge analyses with scheme A (Table 1). As for CladeAge analyses, the percentage of 95% HPD intervals containing the true node age was lower with autocorrelated branch rates, and remained around 95% with uncorrelated branch rates or when MCMC sampling from the prior only. In seven out of nine comparisons, however, the 95% HPD intervals were slightly wider when estimated with the FBD model than with CladeAge scheme A. The FBD model, used with a fixed sampling rate, also appeared somewhat less robust to the violation of the assumed clock model (i.e. with branch-rate autocorrelation), except when the lowest sampling rate *ψ* = 0.01 was used for dataset generation.

In contrast, when the sampling rate was not fixed in FBD analyses, 95% HPD intervals remained similarly wide regardless of the true sampling rate used in dataset generation, and relatively small percentages of 95% HPD intervals contained the true node age in analyses using the posterior (Fig. 3c). A particularly low percentage of 95% HPD intervals (63.5-63.8%; Table 1) contained the true node ages in analyses of datasets generated with autocorrelated branch rates and high or intermediate sampling rates (*ψ* = 0.1 or *ψ* = 0.03). Low accuracy with the FBD model in which sampling rates were not fixed was mostly due to overestimation of intermediate node ages (Fig. 3e, Supplementary Figure S3b-c). The overestimation of node ages in these analyses coincides with a substantial understimation of the sampling rate itself (Supplementary Figure S4). In analyses using the prior alone, the sampling rate was on average estimated as only 30.4, 51.4, and 73.9% of the true sampling rate, when the true sampling rate was *ψ* = 0.1, 0.03, or 0. 01, respectively. Also when sampling from the posterior in analyses of datasets generated with uncorrelated or autocorrelated branch rates, these percentages remained nearly identical (Supplementary Figure S3).

## Applying CladeAge Calibration Densities to Resolve Divergence Times of Cichlid Fishes

### Phylogeography of Cichlidae

Fishes of the percomorph family Cichlidae are known for their extraordinary species richness, which includes the replicated adaptive radiations of the East African Lakes Tanganyika, Malawi, and Victoria (Salzburger et al. 2014). Three reciprocally monophyletic subfamilies occur in Africa and the Middle East (Pseudocrenilabrinae; ∽2100 spp.; see Supplementary Text S2), in South and Central America (Cichlinae; ∽500 spp.), and on Madagascar (Ptychochrominae; 16 spp.). In addition, the most ancestral subfamily Etroplinae consists of two genera, of which *Etroplus* (3 spp.) occurs in Southern India and Sri Lanka while *Paretroplus* (13 spp.) is endemic to Madagascar (Sparks and Smith 2004). As the distribution of cichlids is mostly limited to landmasses of the former supercontinent Gondwana, their biogeography is traditionally considered a product of Gondwanan vicariance (Chakrabarty 2004; Sparks and Smith 2005; Smith et al. 2008; Azuma et al. 2008). In this scenario, the divergence of African Pseudocrenilabrinae and South American Cichlinae must have occurred before or during the break-up of the two continents about 100 Ma (Heine et al. 2013), and Indian and Malagassy members of Etroplinae must have separated before 85 Ma (Ali and Aitchison 2008). Regardless of whether cichlids colonized Africa or South America first, this colonization should have occurred before 120 Ma, as Madagascar and India were separated by that time from both Africa and Antarctica, through which a connection to South America could have existed previously (Ali and Aitchison 2008; Ali and Krause 2011).

However, a Gondwanan history is not supported by the fossil record of Cichlidae. The earliest record of Cichlidae is provided by †*Mahengechromis* spp. from the Mahenge palaeolake, Tanzania (46-45 Ma) (Murray 2000a), followed by the first occurences of neotropical cichlids, recovered from the Argentinian Lumbrara Formation (Malabarba et al. 2006; Alano Perez et al. 2010; Malabarba et al. 2010). The fossiliferous “Faja Verde” layer of the Lumbrera Formation has frequently been cited as Ypresian-Lutetian in age (48.6 Ma) (e.g. Alano Perez et al. 2010); however, it is unclear how the date was obtained (Friedman et al. 2013). The Lumbrera Formation has been assigned to the Casamayoran South American Land Mammal Age (SALMA) (del Papa et al. 2010), which corresponds to 45.4-38.0 Ma (as it is defined by polarities C20-C18) (Vucetich et al. 2007). This age can be further constrained by radiometric dating of a tuff layer overlying the “Faja Verde” deposits, providing a minimum age of 39.9 Ma (del Papa et al. 2010; this evidence seems to have been misinterpreted in a previous analysis of cichlid divergence dates where a minimum age of 33.9 Ma was assumed; Friedman et al. 2013).

Due to the lack of cichlid remains older than 46 Ma, long ghost lineages would need to be postulated in order to reconcile the biogeography of cichlid fishes with Gondwanan vicariance. On the other hand, trans-oceanic dispersal over hundreds or thousands of kilometers, followed by successful colonization of a new continent, appears extremely improbable, given that cichlids are found almost exclusively in freshwater. Whereas several cichlid species occur in brackish-water estuaries and some species are known to tolerate marine saltwater conditions (Myers 1949; Stickney 1986; Uchida et al. 2000), none have ever been observed in the open ocean, more than a few miles from the coast (Conkel 1993; Greenfield and Thomserson 1997). Thus, a long-standing debate has centered on the relative probabilities of the two alternative scenarios, Gondwanan vicariance despite long ghost lineages, or trans-oceanic dispersal despite a common freshwater lifestyle (Vences et al. 2001; Murray 2001a; Chakrabarty 2004; Sparks and Smith 2005; Genner et al. 2007; Smith et al. 2008). However, arguments for both sides have mostly been verbal, and the probabilities of the long ghost lineages required for the Gondwanan vicariance scenario could not properly be quantified, as an objective basis has been lacking for the specification of calibration densities in previous divergence-time analyses (e.g. Azuma et al. 2008; but see Friedman et al. 2013). In contrast, CladeAge calibration densities are based on sampling rate estimates and thus directly account for probabilities of individual ghost lineage durations. In combination with a large-scale molecular phylogeny including multiple cichlid and outgroup fossil constraints, the CladeAge method is therefore ideally suited to assess the most plausible phylogeographic scenario for cichlid fishes.

### A multi-marker phylogeny of teleost fishes

In order to time-calibrate cichlid divergences, we applied CladeAge calibration densities to a large-scale phylogeny of cichlid and outgroup taxa, including nearly 150 fossil constraints. As a first step, we compiled a molecular data set for 40 mitochondrial and nuclear markers, sequenced from 1187 species of the teleost Supercohort Clupeocephala (see Betancur-R et al. 2013). Of the species included in the data set, 578 were members of the family Cichlidae, 516 were members of other families of Cohort Euteleosteomorpha, and 93 species were members of Cohort Otomorpha, the sister lineage of Euteleosteomorpha, and were collectively used as an outgroup in our phylogenetic analysis. Out of a total of 11 050 sequences, 9970 were retrieved from the NCBI nucleotide database (www.ncbi.nlm.nih.gov/nuccore), 85 were obtained from annotated genomes of the Ensembl database (Cunningham et al. 2015), 5 mt-co1 sequences were downloaded from the Barcode of Life Data System (BOLD; Ratnasingham and Hebert 2007), and 328 sequences were identified from other non-annotated genomic resources (Supplementary Tables S2-S7). In addition, 662 sequences of 19 markers were produced specifically for this study, including 26 mitochondrial genomes (see Supplementary Text S1 for sequencing protocols and Supplementary Tables S2 and S5 for accession numbers).

For each marker, sequences were aligned with MAFFT v.7.122b (Katoh and Standley 2013), visually inspected, and poorly aligned regions were removed. Alignments were subsequently divided into primary data blocks according to codon position. In combination, the alignments included 35 817 sites with an overall proportion of undetermined characters of 82.84%. Assuming a general time-reversible model of sequence evolution with gamma-distributed rate variation among sites (GTR+Γ), the fit of partitioning schemes was assessed according to the Bayesian Information Criterion (BIC). The best-fitting partitioning scheme determined with the greedy algorithm implemented in PartitionFinder v.1.0.1 (Lanfear et al. 2012) combined primary data blocks into 30 different partitions (Supplementary Table S8).

Maximum likelihood (ML) phylogenetic tree search was conducted with RAxML v.7.3.1 (Stamatakis 2006; Pfeiffer and Stamatakis 2010), applying unlinked “GTRCAT” models of sequence evolution for each of the 30 partitions. Topological node support was evaluated with RAxML’s rapid bootstrap analysis (option “-f a”) and the “autoMRE” automatic stopping criterion (Stamatakis et al. 2008). Based on the ML phylogeny, we identified 455 clades that were potentially suitable for time calibration, as they were supported by high bootstrap values in our study (≥ 93% with only 6 exceptions) and corroborated by previously published molecular phylogenetic analyses and morphological synapomorphies (Supplementary Figure S5 and Supplementary Text S2). Of the 455 clades, 362 were mutually exclusive and in their sum represented nearly the entire species richness of Clupeocephala (> 99.5%; Supplementary Table S9). This is important in analyses with CladeAge calibration densities, as it ensures that the sister groups of clades used for time calibration are present in the phylogeny, even if their identity is not known prior to the phylogenetic analysis. If sister groups were instead missing from the taxon set, the stem lineages of calibrated clades would appear older than they are, potentially leading to biased age estimates.

### CladeAge model parameter estimation

CladeAge calibration densities are calculated based on estimates of rates of sampling, net diversification, and turnover. In order to use CladeAge calibration densities for the time-calibration of teleost divergences, we obtained estimates for these three parameters from previous studies. Net diversification and turnover rates of teleost fishes were estimated by Santini et al. (2009) as 0.041-0.081 per lineage per million years (L^−1^myr^−1^) and 0.0011-0.37 L^−1^myr^−1^, respectively. These estimates are comparable to those of a more recent analysis by Rabosky et al. (2013), who estimated a mean net diversification rate of 0.098 L^−1^myr^−1^ and a mean turnover rate of 0.284 L^−1^myr^−1^ using a Bayesian model of diversification with rate shifts. We here apply the slightly lower diversification rate estimates of Santini et al. (2009) to calculate CladeAge calibration densities and note that their distributions will tend to be wider, and thus older, than distributions calculated with the rate estimates of Rabosky et al. (2013).

Sampling probabilities have been estimated from the fossil record for a variety of groups and with a wide range of methods. For bony fishes (Osteichthyes) including Clupeocephala, an estimate of the sampling probability was calculated by Foote and Sepkoski (1999) from the frequency ratio *f*_2_^2^/(*f*_1_*f*_3_), where *f*_1_, *f*_2_, and *f*_3_ are the frequencies of genera with stratigraphic ranges of one, two, and three geologic time intervals, respectively (Foote and Raup 1996). The resulting estimate of 0.15-0.30 (Foote and Miller 2007) thus represents the probability that one or more members of a given genus are sampled from a geological time interval, and Foote and Sepkoski (1999) used five-million-year time intervals in their analysis. As CladeAge calibration densities are calculated from instantaneous species-level sampling rates, we translated the genus-level sampling probability estimate of Foote and Sepkoski (1999) as follows. We downloaded the list of all valid scientific names of bony fishes from the Catalogue of Life database (Roskov et al. 2015) and determined the frequency distribution of extant bony fish genus sizes from these names. We then used this distribution in combination with species-level sampling rates to simulate bony fish preservation over five million years and recorded the proportion of genera that were sampled during this interval. The species-level sampling rate was optimized until the resulting proportion of sampled genera was sufficiently close to the genus-level estimate of (Foote and Miller 2007). This optimization was performed separately for the lower and upper bound of estimate of (Foote and Miller 2007). We find that species-level instantaneous sampling rates of 0.0066-0.01806 L^−1^myr^−1^ provide the best fit to five-million-year genus-level preservation probabilities of bony fishes, under the assumption of constant rates and a constant genus-size frequency distribution. For comparison, and in order to provide species-level estimates for future users of CladeAge, we compiled a comprehensive list of published sampling rates in Supplementary Table S1, using the above translation where necessary.

### Divergence-time estimation of teleost fishes

We analyzed the published fossil record for each of the 455 strongly supported teleost clades, and identified their first occurrences, the rock formation in which the earliest record was found, as well as the minimum and maximum age of this formation. Detailed information of the fossil record of each clade is given Supplementary Text S1. According to calibration scheme A, we used first occurrences to define CladeAge calibration densities distributions even if earlier records were known in sister clades, and we reused calibration densities for more inclusive clades if these (i) had no earlier fossil record on their own, but were (ii) either morphologically recognizable or characterized by a discrete geographical distribution so that fossil finds could in principle have been assigned to them directly rather than to parental clades only. For example, the Miocene *Nandopsis* †woodringi represents the earliest record of the genus *Nandopsis,* to which it can be assigned based on the presence of lingual cusps on the oral teeth and four anal-fin spines, a character combination which within cichlids is unique to members of this genus (Chakrabarty 2007). However, *Nandopsis* †woodringi also represents the first occurrence of the clade “SCAC+NCAC”, combining the groups “SCAC” (Southern Central American Clade) and “NCAC” (Northern Central American Clade) of López-Fernández et al. (2010) with a total of 19 genera of Neotropical cichlids. This clade is strongly supported by molecular phylogenies (López-Fernández et al. 2010, this study), but is not characterized by known synapomorphies or a geographical distribution that separates it from its potential sister groups. Thus, if stem-fossils were found of clade “SCAC+NCAC”, these would likely be misassigned to the next more inclusive clade that is morphologically recognizable, in this case the tribe Heroini. A lack of recognizable features for a clade thus effectively reduces the sampling rate of its stem lineage to 0. In order to account for this reduction, CladeAge calibration densities were defined exclusively for morphologically (or in some cases geographically) recognizable clades. We identified fossil constraints for a total of 147 clades, including 18 clades within cichlids (see Supplementary Text S1 and Supplementary Figure S5).

In order to reduce model complexity and increase compuational efficiency of Bayesian phylogenetic inference, eight markers with the greatest proportions of missing sequences were removed from the data set (Supplementary Table S4). In addition, a total of 80 codon positions with signatures of episodic selection were identified with the mixed effects model of evolution implemented in HyPhy (Murrell et al. 2012; Kosakovsky Pond et al. 2005) and removed from the alignment. We further collapsed each of the 362 mutually exclusive clades to individual tips, and for each marker we chose sequences of clade members at random to represent the terminal clade. To account for sequence variation within a clade, we repeated random sequence sampling five times, producing five replicate datasets that each included a total of 27 950 sites with 59.3% missing data. Each of the five replicate data sets was used for phylogenetic inference and time calibration with BEAST, on the basis of 147 CladeAge calibration densities. As for ML analyses, the data set was partitioned according to marker and codon position. Tree topology and branch lengths were linked among partitions, but parameters of the clock and sequence substitution models remained unlinked. We assumed an uncorrelated relaxed molecular clock (Drummond et al. 2006) and applied the reversible-jump based substitution model of Bouckaert et al. (2013). For each partition, a gamma distribution of among-site rate heterogeneity with four rate categories was assumed. We used the flexible birth-death skyline model (Stadler et al. 2012) with independent diversification rate parameters for the pre-Cretaceous Mesozoic (> 145.5 Ma), the Early (145.5-99.6 Ma) and Late Cretaceous (99.6-66.0 Ma), as well as the Cenozoic (< 66.0 Ma), and specified a sampling fraction *ρ* of 0.0135 according to the ratio of tips included in the analysis to the total extant diversity of Clupeocephala. We left the tree topologically unconstrained except for nodes used for time calibration. Justifications for the assumed monophyly of each clade used for time-calibration are given in Supplementary Text S2. For each data set replicate, 600 million MCMC states were sampled, and we repeated the analysis without data, sampling from the prior to ensure that conclusions were not pre-determined by the prior.

### Resulting timeline of cichlid and teleost divergences

Comparison of MCMC traces between run replicates suggested that all replicates had converged at the same posterior distribution. After discarding 60 million MCMC generation of each replicate run as burn-in, we produced a joint posterior tree sample with 1000 trees per replicate, and generated a Maximum Clade Credibility (MCC) tree from the combined distribution of 5000 posterior trees. The inferred timeline of cichlid and outgroup teleost divergences is summarized in Figure 4a and shown in more detail in Supplementary Figure S6.

**Figure 4:**
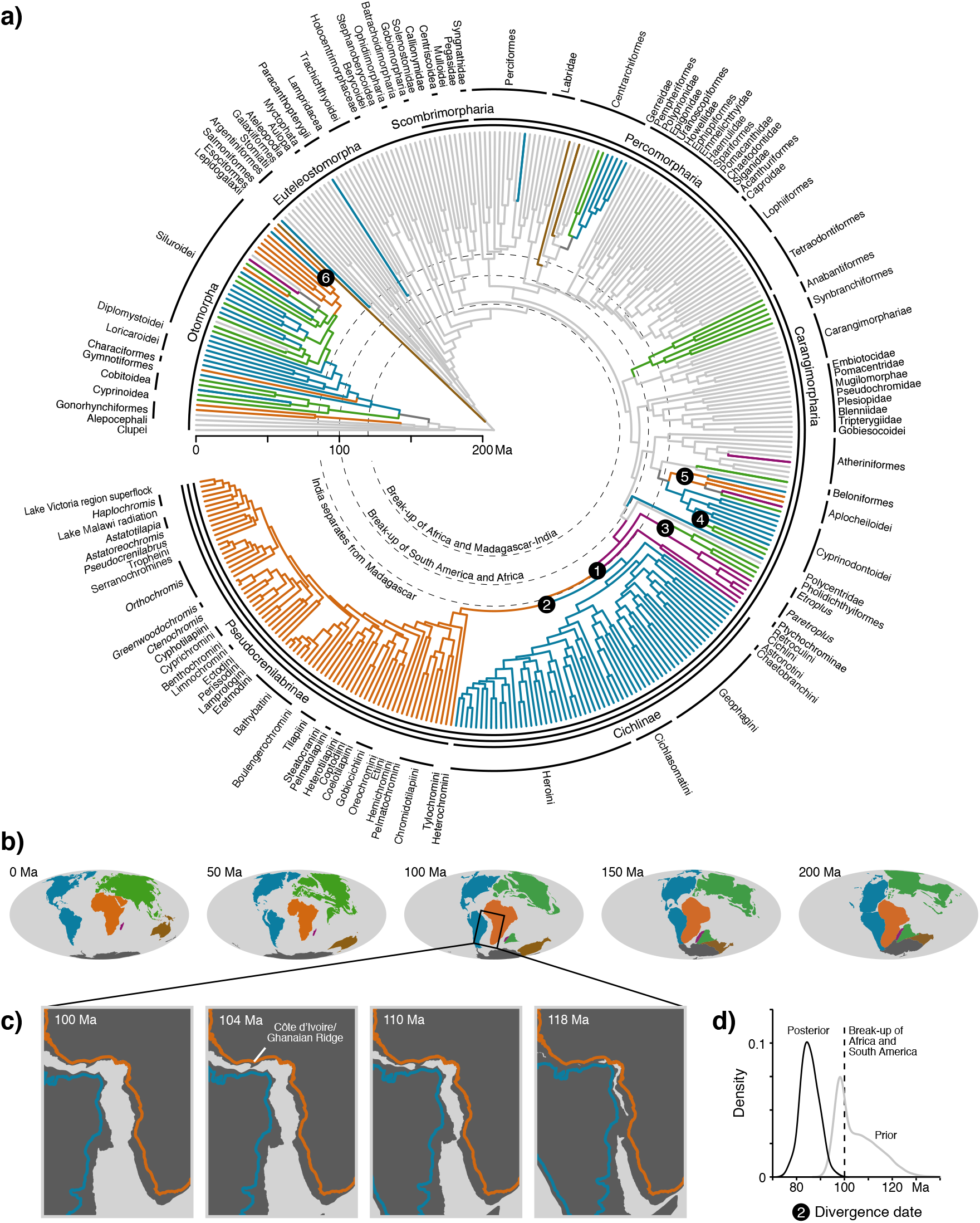
Time-calibrated phylogenetic tree of teleost fishes and plate tectonic reconstructions. a) Maximum Clade Credibility phylogeny of cichlid and outgroup teleost fishes, time-calibrated with 147 fossil constraints. Dashed lines mark continental break-up events of Gondwanan landmasses. Colors of terminal branches indicate the center of diversity for clades occurring exclusively in freshwater or brackish water habitats (blue: America; orange: Africa; purple: Madagascar; brown: Australia; green: India and Eurasia). Groups with marine representatives are shown in gray. Colors of internal branches indicate past distributions according to the most parsimonious scenario of dispersal and freshwater colonization, taking into account past geographic distances between landmasses. Dark gray branches indicate equal parsimony of multiple scenarios. Six dispersal events with particularly strong evidence for trans-oceanic dispersal are highlighted: 1) Since the two oldest cichlid subfamilies, Etroplinae and Ptychochrominae, occur on Madagascar (and Ptychochrominae being endemic to Madagascar), this landmass represents the most likely origin of family Cichlidae. According to our timeline of teleost divergences, dispersal of the clade combining the younger two subfamilies Pseudocrenilabrinae and Cichlinae from Madagascar to either Africa or South America occurred after 85.7 Ma (95% HPD: 93.8-77.8 Ma), substantially later than the latest possible separation of Madagascar from both landmasses around 120 Ma (Ali and Aitchison 2008; Ali and Krause 2011). Since Madagascar was geographically closer to Africa than to South America at 85.7 Ma, we assume that cichlids dispersed to Africa before reaching South America. 2) The divergence event of African Pseudocrenilabrinae and South American Cichlinae is estimated at 81.8 Ma (95% HPD: 89.4-74.0 Ma), long after the final separation of the two continents at 104-100 Ma Heine et al. (2013). 3) Within the cichlid subfamily Etroplinae, the Indian genus *Etroplus* and the Malagassy genus *Paretroplus* diverged about 69.5 Ma (95% HPD: 85.9-53.1 Ma), probably after the break-up of India and Madagascar between 90-85 Ma (Ali and Aitchison 2008). 4) The predominantly American Cyprinodontoidei include multiple Old World lineages, such as the clade combining the Mediterranean *Aphanius* and Valenciidae, which diverged from South American relatives about 50.6 Ma (95% HPD: 61.4-39.3 Ma). 5) With an estimated crown age 80.8 Ma (95% HPD: 92.5-69.7 Ma), the cyprinodontiform suborder Aplocheiloidei includes American, African, Malagassy, and Indian lineages of strict freshwater fishes. The aplocheilid sister genera *Pachypanchax* and *Aplocheilus* occur in Madagascar and Asia, respectively, and diverged about 42.8 Ma (95% HPD: 60.4-23.8 Ma). 6) The Mexican *Lacantunia enigmatica* appears deeply nested within African freshwater Siluroidei, but separated about 49.6 Ma (95% HPD: 57.9-45.2 Ma). b) Plate tectonic reconstructions of the break-up of Gondwana between 200 Ma and the present. c) Stages in the separation of South America and Africa between 118 and 100 Ma. According to the plate kinematic model of Heine et al. (2013), final breakup in the South Atlantic Rift System (SARS) occured between 113-112 Ma in the outer Santos Basin. African and South American lithospheres completely separated at 104 Ma, whereby the last continental connection remained along the Côte d’Ivoire/Ghanaian Ridge in the Equatorial Rift System (EqRS). Orange and blue outlines represent Africa and South America with present coastlines. Dark gray shapes indicate the restored continental margin (see Heine et al. 2013). Modified from Heine et al. (2013). d) Prior and posterior distributions for the divergence date of African and South American cichlid fishes (marked with “2” in a). 99.93% of the posterior probability mass supports a divergence event younger than 100 Ma, and thus trans-Atlantic dispersal instead of Gondwanan vicariance. In contrast, this scenario is supprted by only 65.6% of the prior probability.

The MCC tree topology was well supported, and corroborates the higher-level groupings found in recent large-scale Bayesian phylogenies of teleost fishes (Near et al. 2013; Betancur-R et al. 2013), as well as previously identified relationships within cichlid fishes (e.g. Schwarzer et al. 2009; López-Fernández et al. 2010; Meyer et al. 2015). With a single exception (Centropomidae), all unconstrained clades from our list of 455 clades were recovered as monophyletic. According to the MCC timeline, crown Clupeocephala originated around 207.8 Ma (95% HPD: 234.5-186.2 Ma), crown Acanthomorphata appeared 144.2 Ma (95% HPD: 158.4-130.6 Ma), and South American Cichlinae and African Pseudocrenilabrinae diverged about 81.6 Ma (95% HPD: 89.4-74.0 Ma). In comparison, the age of crown Clupeocephala appears markedly older in the studies of Betancur-R et al. (2013) and Near et al. (2013), who estimated their origin at about 251.1 (95% HPD: 276.1-226.1 Ma) and 273.7 Ma (95% HPD: 307.5-242.0 Ma), respectively. The divergence date of Acanthomorphata is more comparable between the three studies, and was estimated at 164.9 Ma (95% HPD 186.0-144.4 Ma) in Betancur-R et al. (2013), and around 142.5 Ma (95% HPD: 154.0-132.0 Ma) in Near et al. (2013), less than two million years younger than the estimate resulting from our time-calibrated phylogeny. For relatively younger divergences, however, our estimates appear older than those of Betancur-R et al. (2013) and Near et al. (2013): The divergence of South American Cichlinae and African Pseudocrenilabrinae was estimated at 62.0 Ma (95% HPD: 70.4-53.9 Ma) in Betancur-R et al. (2013) and as young as 26.0 Ma (95% HPD: 29.6-22.0 Ma) in Near et al. (2013). Notably even the older limit of the 95% HPD of the latter estimate is predated by at least 5 well-characterized fossil species within crown Pseudocrenilabrinae (Murray 2001b) and crown Cichlinae (Malabarba et al. 2010, 2006; Alano Perez et al. 2010; Malabarba and Malabarba 2008) and thus is in strong disagreement with the cichlid fossil record. Thus, the consistent application of CladeAge divergence date probability distributions to all clades with known fossil records appears to remove conflicts of comparatively younger node ages with the fossil record, while at the same time reducing the ghost lineages for relatively older clades. A more detailed comparison of clade age estimates between the three studies is shown in Supplementary Figure S7.

While our age estimates for cichlid divergences are generally older than those obtained in Betancur-R et al. (2013) and Near et al. (2013), they are still markedly too young to support strictly Gondwanan vicariance between Indian, Malagassy, Neotropical, and African groups of cichlid fishes, as well as within other groups of freshwater fishes included in our phylogeny (Fig. 4). Notably, the divergence of Neotropical Cichlinae and African Pseudocrenilabrinae, estimated at 81.6 Ma, appears to have occurred about 20 myr after the final separation of the American and African landmasses at 104-100 Ma Heine et al. (2013).

Comparison of these results with those obtained by MCMC sampling from the prior distribution shows that the divergence estimate for African and South American cichlids is driven by the molecular sequence data. The prior distribution is markedly older than the divergence date posterior for this split, with 65.6% of the prior samples being younger than 100 Ma, whereas the same is true for 99.93% of the posterior distribution (Fig. 4d). As a consequence, the Bayes factor in favour of a divergence younger than 100 Ma is 752, which can be considered overwhelming evidence (Kass and Raftery 1995) supporting the trans-Atlantic dispersal scenario, as opposed to Gondwanan vicariance.

## Discussion

### Divergence-time estimation with CladeAge calibration densities

Our analyses of datasets simulated under a wide range of conditions show that CladeAge calibration densities allow bias-free estimation of divergence times. The comparison of four different calibration schemes confirms that calibration scheme A (Fig. 2a) performs better than other schemes and should thus be applied whenever CladeAge calibration densities are used for time calibration. This implies that for each clade, the oldest fossil record of this clade should be used as a time constraint, regardless of whether the fossil is also the earliest record of other (nested or parental) clades, and even if the fossil is younger than the oldest fossil record of the clade’s sister group. With calibration scheme A, the CladeAge model produces very similar results to the FBD model with fixed sampling rates, but appears slightly more robust to model violation in the form of branch-rate autocorrelation, at least with larger numbers of fossil calibrations. When the sampling rate is not fixed in FBD analyses and only the oldest fossils of each clade are used for time calibration, the sampling rate is often substantially underestimated, leading to wide confidence intervals and overestimation of node ages. This suggests that when the sampling-rate parameter is not fixed in analyses with the FBD model, a rather complete representation of the fossil record should be included in the analysis instead of only the oldest fossil of each clade.

However, for the practical time calibration of higher-level phylogenies, the compilation of the entire fossil record of clades used for calibration may be far less feasible than the identification of their oldest reported fossils. A large amount of paleontological literature has been dedicated to determine oldest taxon appearances across the tree of life, and demonstrates the difficulties associated with the identification of these records as well as of their ages (Benton 1993; Benton and Donoghue 2007; Hedges and Kumar 2009; Ksepka et al. 2011; Benton et al. 2015). While the Paleobiological Database (www.paleobiodb.org) provides information not only about the oldest fossils of clades, but about a much larger number of fossils for many clades, the taxonomic assignment and age ranges given for these fossils are usually far less well curated than those of first taxon appearances that are dealt with in dedicated literature. In addition, neither the paleontological literature nor databases that use information from this literature are likely to provide an unbiased representation of the age distribution of fossils within a clade. Instead, new discoveries of fossils that extend the known age range of clades are almost guaranteed to be reported in the literature (and as a consequence also in databases), whereas younger findings may often not be considered worthy of publication. Thus, available information about the oldest records of clades is likely to be better curated and less biased than the collective data for all its fossils. Furthermore, since fossils are added as tips in analyses with the FBD model, the computational demand increases with the number of fosssils, and may be prohibitive for higher-level phylogenies of clades with rich fossil records.

On the other hand, the specification of CladeAge calibration densities is computationally not more demanding than any other calibration density used in node dating, and is thus suitable for large-scale phylogenetic analyses. In contrast to the current software implementations of the FBD model (Gavryushkina et al. 2014; Zhang et al. 2015), the CladeAge method can also account for different rates of diversification and sampling in different co-existing clades, as calibration densities are independently specified for each calibrated clade. This is likely to improve age estimates in higher-level phylogenies such as the vertebrate tree of life, where substantial differences in these rates have previously been demonstrated (Foote and Sepkoski 1999; Alfaro et al. 2009). Thus, a strategy for the Bayesian divergence time estimation of large trees like the vertebrate tree of life could include the following steps. First, representative groups with suitable fossil records could be chosen from several of the higher taxa (i.e. mammals, birds, teleost fishes) included in the phylogeny, and could be used to estimate sampling and diversification rate parameters for these taxa. This could be done either using information from the fossil record alone (Silvestro et al. 2014; Starrfelt and Liow 2015), or in combination with molecular data, e.g. by means of separate FBD analyses for each of the representative groups. Then, the resulting rate estimates could be used to calibrate the ages of clades within the higher taxa, under the assumption that the true rates of these clades are comparable to those estimated from representative groups. Finally, divergence time estimation of the complete phylogeny could be carried out with node dating, on the basis of CladeAge calibration densities.

### Trans-Atlantic dispersal of cichlid fishes

Using CladeAge calibration denstities for 147 clades of teleost fishes, we found strong evidence for trans-oceanic dispersal, not only in cichlid fishes, but also in several other groups of freshwater fishes, including Cyprinodontoidei, Aplocheiloidei, and Siluroidei (Fig. 4). The calibration densities used in our analysis were based on estimates of sampling and diversification rates by Foote and Sepkoski (1999) and Santini et al. (2009), and our results could thus be biased if these estimates are inaccurate. We note, however, that the rate estimates used by us are low compared to those of other clades (see Supplementary Table S1) or those obtained by other authors (Rabosky et al. 2013). Thus, the used estimates for sampling and diversification rates are more likely underestimates than overestimates, which would lead to calibration densities that are wider than they should be, and therefore to overestimated ages of clades in our phylogeny.

For several further reasons, we would expect our age estimates to be rather over-than underestimated. First, our simulations have shown that age estimates obtained with CladeAge calibration densities (or the FBD model) can appear too old when the sampling rate is low and the assumed clock model is violated, e.g. by branch-rate autocorrelation (Supplementary Figure S3b). In practice, autocorrelation of branch-specific substitution rates can rarely be excluded, and may be present also in teleost fishes, as many factors influencing rate variation are heritable in vertebrates (Nabholz et al. 2008; Amster and Sella 2016). Second, while our molecular dataset is composed of both nuclear and mitochondrial sequences, nuclear sequences were available to a greater degree for taxa outside of Cichlidae, and may be underrepresented for clades within this family. As the substitution rate of mitochondrial markers is usually higher than that of nuclear markers (Brown et al. 1979), overall genetic divergences between cichlids might appear higher than those of other groups that have a similar age but a lower proportion of missing data in nuclear markers. Third, by using concatenation of all sequence markers rather than the multispecies coalescent-model (which would have been computationally infeasible), we essentially ignored potential variation between gene trees due to incomplete lineage sorting, which has been shown to lead to inflated age estimates in several studies (McCormack et al. 2011; Colombo et al. 2015). Fourth, in contrast to the authors of a previous study on cichlid divergence times (Friedman et al. 2013), we assumed nested positions of the oldest Neotropical and African cichild fossils within the subfamilies Cichlinae and Pseudocrenilabrinae, respectively, rather than positions along their stem lineages. Specifically, we assumed a position within genus *Gymnogeophagus* for *Gymnogeophagus* † *eocenicus,* a position of † *Tremembichthys garciae* within Cichlasomatini, a position of † *Plesioheros chauliodus* within Heroini, a position of †*Proterocara argentina* within a clade formed by the extant genera *Teleocichla* and *Crenicichla,* and a position of † *Mahengechromis* spp. within the African tribe Hemichromini, which are all supported by morphological analyses (Murray 2000b, 2001b; Malabarba and Malabarba 2008; Smith et al. 2008; Alano Perez et al. 2010; Malabarba et al. 2010). If these nested positions should be unjustified (as suggested by Friedman et al. 2013), even younger ages of cichlid divergences would be expected. Taken together, our analyses strongly support trans-Atlantic dispersal of cichlids, between 89.4 and 74.0 Ma, or earlier.

## Conclusion

In this study, we have presented a new approach to Bayesian node dating that is directly based on probabilities of fossil sampling, and thus overcomes previous shortcomings of node dating. We have demonstrated that our approach allows accurate and precise divergence-time estimation and represents a viable alternative to the FBD model when estimates for the rates of fossil sampling and diversification are available *a priori.* Our approach is particularly suitable for the time calibration of large-scale phylogenies, and we have outlined strategies how to use our method in order to account for variable rates of sampling and diversification in different clades. By applying our approach to a detailed phylogeny of teleost fishes, we have shown that freshwater fishes in several clades have diverged long after the separation of the continents on which they live, which implies that fishes from these clades have successfully traversed oceanic environments despite their adaptations to a freshwater lifestyle. These examples include the trans-Atlantic dispersal of cichlid fishes, which led to their colonization of South and Central American rivers and lakes, and to the radiation of Neotropical cichlid fishes into over 600 extant species. We have implemented our approach in the CladeAge package for BEAST (Gavryushkina et al. 2014; Bouckaert et al. 2014), which is freely available at www.beast2.org.

## Acknowledgements

We would like to thank Alison Murray and Ken Monsch for advice on fossil calibrations, and Jaroslav Vohánka, Jérémy Winninger and Brigitte Aeschbach for help during labwork and 454 pyrosequencing. Alexei Drummond, Sebastian Höhna, Hallie Sims, Tanja Stadler, and Christian Heine provided valuable comments on the project and the manuscript. Discussions with Mike Foote and David Bapst stimulated the translation of genus-level to species-level sampling rates. We wish to acknowledge the contribution of the NeSI (www.nesi.org.nz) and Abel (www.hpc.uio.no) high-performance computing facilities to the results of this research.

## References

Alano Perez, P., M. C. Malabarba, and C. Del Papa. 2010. A new genus and species of Heroini (Perciformes: Cichlidae) from the early Eocene of southern South America. Neotrop. Ichthyol. 8:631–642.

Alfaro, M. E., F. Santini, C. D. Brock, H. Alamillo, A. Dornburg, D. L. Rabosky, G. Carnevale, and L. J. Harmon. 2009. Nine exceptional radiations plus high turnover explain species diversity in jawed vertebrates. Proc. Natl. Acad. Sci. USA 106:13410–13414.

Ali, J. R. and J. C. Aitchison. 2008. Gondwana to Asia: Plate tectonics, paleogeography and the biological connectivity of the Indian sub-continent from the Middle Jurassic through latest Eocene (166-35 Ma). Earth-Sci. Rev. 88:145–166.

Ali, J. R. and D. W. Krause. 2011. Late Cretaceous bioconnections between Indo-Madagascar and Antarctica: refutation of the Gunnerus Ridge causeway hypothesis. J. Biogeogr. 38:1855–1872.

Amster, G. and G. Sella. 2016. Life history effects on the molecular clock of autosomes and sex chromosomes. Proc. Natl. Acad. Sci. USA Early Access, doi: 10.1073/pnas.1515798113.

Arcila, D., R. A. Pyron, J. C. Tyler, G. Ortí, and R. Betancur-R. 2015. An evaluation of fossil tip-dating versus node-age calibrations in tetraodontiform fishes (Teleostei: Percomorphaceae). Mol. Phylogenet. Evol. 82:131–145.

Azuma, Y., Y. Kumazawa, M. Miya, K. Mabuchi, and M. Nishida. 2008. Mitogenomic evaluation of the historical biogeography of cichlids toward reliable dating of teleostean divergences. BMC Evol. Biol. 8:215.

Beaulieu, J. M. and M. J. Donoghue. 2013. Fruit evolution and diversification in campanulid angiosperms. Evolution 67:3132–3144.

Beck, R. M. D. and M. S. Y. Lee. 2014. Ancient dates or accelerated rates? Morphological clocks and the antiquity of placental mammals. Proc. R. Soc. B 281:20141278–20141278.

Benton, M. and P. Donoghue. 2007. Paleontological evidence to date the tree of life. Mol. Biol. Evol. 24:26–53.

Benton, M. J., ed. 1993. The fossil record 2. Chapman & Hall, London, UK.

Benton, M. J., M. J. Donoghue, R. J. Asher, M. Friedman, T. J. Near, and J. Vinther. 2015. Constraints on the timescale of animal evolutionary history. Palaeontol. Electron. 18.1.1FC:1–106.

Betancur-R, R., R. E. Broughton, E. O. Wiley, K. E. Carpenter, J. A. López, C. Li, N. I. Holcroft, D. Arcila, M. D. Sanciangco, J. C. Cureton, F. Zhang, T. Buser, M. A. Campbell, J. A. Ballesteros, A. Roa-Varon, S. C. Willis, W. C. Borden, T. Rowley, P. C. Reneau, D. J. Hough, G. Lu, T. Grande, G. Arratia, and G. Ortí. 2013. The Tree of Life and a new classification of bony fishes. PLOS Currents: Tree of Life Pages 1–45.

Bouckaert, R., M. V. Alvarado-Mora, and J. R. Rebello Pinho. 2013. Evolutionary rates and HBV: issues of rate estimation with Bayesian molecular methods. Antivir. Ther. 18:497–503.

Bouckaert, R., J. Heled, D. Kühnert, T. Vaughan, C.-H. Wu, D. Xie, M. A. Suchard, A. Rambaut, and A. J. Drummond. 2014. BEAST 2: a software platform for Bayesian evolutionary analysis. PLOS Comput. Biol. 10:e1003537.

Bromham, L. and D. Penny. 2003. The modern molecular clock. Nat. Rev. Genet. 4:216–224.

Brown, W. M., M. J. George, and A. C. Wilson. 1979. Rapid evolution of animal mitochondrial DNA. Proc. Natl. Acad. Sci. USA 76:1967–1971.

Chakrabarty, P. 2004. Cichlid biogeography: comment and review. Fish Fish. 5:97–119.

Chakrabarty, P. 2007. Taxonomic status of the hispaniolan Cichlidae. Occas. Pap. Mus. Zool. Univ. Mich. Pages 1–20.

Claramunt, S. and J. Cracraft. 2015. A new time tree reveals Earth history’s imprint on the evolution of modern birds. Sci. Adv. 1:e1501005.

Colombo, M., M. Damerau, R. Hanel, W. Salzburger, and M. Matschiner. 2015. Diversity and disparity through time in the adaptive radiation of Antarctic notothenioid fishes. J. Evol. Biol. 28:376–394.

Conkel, D. 1993. Cichlids of North and Central America. TFH Publications, Neptune City, New Jersey.

Cunningham, F., M. R. Amode, D. Barrell, K. Beal, K. Billis, S. Brent, D. Carvalho-Silva, P. Clapham, G. Coates, S. Fitzgerald, L. Gil, C. G. Giron, L. Gordon, T. Hourlier, S. E. Hunt, S. H. Janacek, N. Johnson, T. Juettemann, A. K. Kahari, S. Keenan, F. J. Martin, T. Maurel, W. McLaren, D. N. Murphy, R. Nag, B. Overduin, A. Parker, M. Patricio, E. Perry, M. Pignatelli, H. S. Riat, D. Sheppard, K. Taylor, A. Thormann, A. Vullo, S. P. Wilder, A. Zadissa, B. L. Aken, E. Birney, J. Harrow, R. Kinsella, M. Muffato, M. Ruiner, S. M. J. Searle, G. Spudich, S. J. Trevanion, A. Yates, D. R. Zerbino, and P. Flicek. 2015. Ensembl 2015. Nucleic Acids Res. 43:D662–D669.

del Papa, C., A. Kirschbaum, J. Powell, A. Brod, F. Hongn, and M. Pimentel. 2010. Sedimentological, geochemical and paleontological insights applied to continental omission surfaces: A new approach for reconstructing an Eocene foreland basin in NW Argentina. J. South Am. Earth Sci. 29:327–345.

dos Reis, M., P. Donoghue, and Z. Yang. 2014. Neither phylogenomic nor palaeontological data support a Palaeogene origin of placental mammals. Biol. Lett. 10:20131003–20131003.

Drummond, A. J., S. Y. W. Ho, M. J. Philips, and A. Rambaut. 2006. Relaxed phylogenetics and dating with confidence. PLOS Biol. 4:e88.

Drummond, A. J., O. G. Pybus, A. Rambaut, R. Forsberg, and A. G. Rodrigo. 2003. Measurably evolving populations. Trends Ecol. Evol. 18:481–488.

Faria, N. R., A. Rambaut, M. A. Suchard, G. Baele, T. Bedford, M. J. Ward, A. J. Tatem, J. D. Sousa, N. Arinaminpathy, J. Pépin, D. Posada, M. Peeters, O. G. Pybus, and P. Lemey. 2014. The early spread and epidemic ignition of HIV-1 in human populations. Science 346:56–61.

Foote, M., J. P. Hunter, C. M. Janis, and J. J. Sepkoski, Jr. 1999. Evolutionary and preservational constraints on origins of biologic groups: divergence times of eutherian mammals. Science 283:1310–1314.

Foote, M. and A. I. Miller. 2007. Principles of Paleontology. 3 ed. W. H. Freeman, New York.

Foote, M. and D. M. Raup. 1996. Fossil preservation and the stratigraphic ranges of taxa. Paleobiology 22:121–140.

Foote, M. and J. J. Sepkoski, Jr. 1999. Absolute measures of the completeness of the fossil record. Nature 398:415–417.

Forest, F. 2009. Calibrating the Tree of Life: fossils, molecules and evolutionary timescales. Ann. Bot. 104:789–794.

Friedman, M., B. P. Keck, A. Dornburg, R. I. Eytan, C. H. Martin, C. D. Hulsey, P. C. Wainwright, and T. J. Near. 2013. Molecular and fossil evidence place the origin of cichlid fishes long after Gondwanan rifting. Proc. R. Soc. B 280:20131733.

Gavryushkina, A., T. A. Heath, D. T. Ksepka, T. Stadler, D. Welch, and A. J. Drummond. 2015. Bayesian total evidence dating reveals the recent crown radiation of penguins. arXiv preprint Pages 1–65.

Gavryushkina, A., D. Welch, T. Stadler, and A. J. Drummond. 2014. Bayesian inference of sampled ancestor trees for epidemiology and fossil calibration. PLOS Comput. Biol. 10:e1003919.

Genner, M. J., O. Seehausen, D. H. Lunt, D. A. Joyce, P. W. Shaw, G. R. Carvalho, and G. F. Turner. 2007. Age of cichlids: new dates for ancient lake fish radiations. Mol. Biol. Evol. 24:1269–1282.

Gernhard, T. 2008. The conditioned reconstructed process. J. Theor. Biol. 253:769–778.

Gire, S. K., A. Goba, K. G. Andersen, R. S. G. Sealfon, D. J. Park, L. Kanneh, S. Jalloh, M. Momoh, M. Fullah, G. Dudas, S. Wohl, L. M. Moses, N. L. Yozwiak, S. Winnicki, C. B. Matranga, C. M. Malboeuf, J. Qu, A. D. Gladden, S. F. Schaffner, X. Yang, P.-P. Jiang, M. Nekoui, A. Colubri, M. R. Coomber, M. Fonnie, A. Moigboi, M. Gbakie, F. K. Kamara, V. Tucker, E. Konuwa, S. Saffa, J. Sellu, A. A. Jalloh, A. Kovoma, J. Koninga, I. Mustapha, K. Kargbo, M. Foday, M. Yillah, F. Kanneh, W. Robert, J. L. B. Massally, S. B. Chapman, J. Bochicchio, C. Murphy, C. Nusbaum, S. Young, B. W. Birren, D. S. Grant, J. S. Scheiffelin, E. S. Lander, C. Happi, S. M. Gevao, A. Gnirke, A. Rambaut, R. F. Garry, S. H. Khan, and P. C. Sabeti. 2014. Genomic surveillance elucidates Ebola virus origin and transmission during the 2014 outbreak. Science 345:1369–1372.

Greenfield, D. W. and J. E. Thomserson. 1997. Fishes of the Continental Waters of Belize. University Press of Florida, Gainesville, Florida.

Grimm, G. W., P. Kapli, B. Bomfleur, S. McLoughlin, and S. S. Renner. 2015. Using more than the oldest fossils: dating Osmundaceae with three Bayesian clock approaches. Syst. Biol. 64:396–405.

Heath, T. A., J. P. Huelsenbeck, and T. Stadler. 2014. The fossilized birth-death process for coherent calibration of divergence-time estimates. Proc. Natl. Acad. Sci. USA 111:E2957–E2966.

Hedges, S. B. and S. Kumar, eds. 2009. The Timetree of Life. Oxford University Press, Oxford, UK.

Heine, C., J. Zoethout, and R. D. Müller. 2013. Kinematics of the South Atlantic rift. Solid Earth 4:215–253.

Heled, J. and A. J. Drummond. 2012. Calibrated tree priors for relaxed phylogenetics and divergence time estimation. Syst. Biol. 61:138–149.

Heled, J. and A. J. Drummond. 2015. Calibrated birth-death phylogenetic time-tree priors for Bayesian inference. Syst. Biol. 64:369–383.

Ho, S. Y. W. and M. J. Phillips. 2009. Accounting for calibration uncertainty in phylogenetic estimation of evolutionary divergence times. Syst. Biol. 58:367–380.

Höhna, S., T. Stadler, F. Ronquist, and T. Britton. 2011. Inferring speciation and extinction rates under different sampling schemes. Mol. Biol. Evol. 28:2577–2589.

Huelsenbeck, J. P. and F. Ronquist. 2001. MRBAYES: Bayesian inference of phylogenetic trees. Bioinformatics 17:754–755.

Jarvis, E. D., S. Mirarab, A. J. Aberer, B. Li, P. Houde, C. Li, S. Y. W. Ho, B. C. Faircloth, B. Nabholz, J. T. Howard, A. Suh, C. C. Weber, R. R. da Fonseca, J. Li, F. Zhang, H. Li, L. Zhou, N. Narula, L. Liu, G. Ganapathy, B. Boussau, M. S. Bayzid, V. Zavidovych, S. Subramanian, T. Gabaldón, S. Capella-Gutiérrez, J. Huerta-Cepas, B. Rekepalli, K. Munch, M. Schierup, B. Lindow, W. C. Warren, D. Ray, R. E. Green, M. W. Bruford, X. Zhan, A. Dixon, S. Li, N. Li, Y. Huang, E. P. Derryberry, M. F. Bertelsen, F. H. Sheldon, R. T. Brumfield, C. V. Mello, P. V. Lovell, M. Wirthlin, M. P. C. Schneider, F. Prosdocimi, J. A. Samaniego, A. M. Vargas Velazquez, A. Alfaro-Núñez, P. F. Campos, B. Petersen, T. Sicheritz-Ponten, A. Pas, T. Bailey, P. Scofield, M. Bunce, D. M. Lambert, Q. Zhou, P. Perelman, A. C. Driskell, B. Shapiro, Z. Xiong, Y. Zeng, S. Liu, Z. Li, B. Liu, K. Wu, J. Xiao, X. Yinqi, Q. Zheng, Y. Zhang, H. Yang, J. Wang, L. Smeds, F. E. Rheindt, M. Braun, J. Fjeldsa, L. Orlando, F. K. Barker, K. A. Jønsson, W. Johnson, K.-P. Koepfli, S. O’Brien, D. Haussler, O. A. Ryder, C. Rahbek, E. Willerslev, G. R. Graves, T. C. Glenn, J. McCormack, D. Burt, H. Ellegren, P. Alström, S. V. Edwards, A. Stamatakis, D. P. Mindell, J. Cracraft, E. L. Braun, T. Warnow, W. Jun, M. T. P. Gilbert, and G. Zhang. 2014. Whole-genome analyses resolve early branches in the tree of life of modern birds. Science 346:1320–1331.

Jetz, W., G. H. Thomas, J. B. Joy, K. Hartmann, and A. Ø. Mooers. 2012. The global diversity of birds in space and time. Nature 491:444–448.

Kass, R. E. and A. E. Raftery. 1995. Bayes Factors. Journal of the American Statistical Association 90:773–795.

Katoh, K. and D. M. Standley. 2013. MAFFT multiple sequence alignment software version 7: improvements in performance and usability. Mol. Biol. Evol. 30:772–780.

Klopfstein, S., L. Vilhelmsen, and F. Ronquist. 2015. A nonstationary Markov model detects directional evolution in hymenopteran morphology. Syst. Biol. 64:1089–1103.

Kosakovsky Pond, S. L., S. D. W. Frost, and S. V. Muse. 2005. HyPhy: hypothesis testing using phylogenies. Bioinformatics 21:676–679.

Kosiol, C., I. Holmes, and N. Goldman. 2007. An empirical codon model for protein sequence evolution. Mol. Biol. Evol. 24:1464–1479.

Ksepka, D. T., M. J. Benton, M. T. Carrano, M. A. Gandolfo, J. J. Head, E. J. Hermsen, W. G. Joyce, K. S. Lamm, J. S. L. Patané, M. J. Phillips, P. D. Polly, M. Van Tuinen, J. L. Ware, R. C. M. Warnock, and J. F. Parham. 2011. Synthesizing and databasing fossil calibrations: divergence dating and beyond. Biol. Lett. 7:801–803.

Lanfear, R., B. Calcott, S. Y. W. Ho, and S. Guindon. 2012. PartitionFinder: combined selection of partitioning schemes and substitution models for phylogenetic analyses. Mol. Biol. Evol. 29:1695–1701.

Lepage, T., D. Bryant, H. Philippe, and N. Lartillot. 2007. A general comparison of relaxed molecular clock models. Mol. Biol. Evol. 24:2669–2680.

Lepage, T., S. Lawi, P. Tupper, and D. Bryant. 2006. Continuous and tractable models for the variation of evolutionary rates. Math. Biosci. 199:216–233.

Lewis, P. O. 2001. A likelihood approach to estimating phylogeny from discrete morphological character data. Syst. Biol. 50:913–925.

López-Fernández, H., K. O. Winemiller, and R. L. Honeycutt. 2010. Multilocus phylogeny and rapid radiations in Neotropical cichlid fishes (Perciformes: Cichlidae: Cichlinae). Mol. Phylogenet. Evol. 55:1070–1086.

Malabarba, M. C. and L. R. Malabarba. 2008. A new cichlid Tremembichthys garcia (Actinopterygii, Perciformes) from the Eocene-Oligocene of Eastern Brazil. Rev. Bras. Paleontol. 11:59–68.

Malabarba, M. C., L. R. Malabarba, and C. Del Papa. 2010. Gymnogeophagus eocenicus, n. sp (Perciformes: Cichlidae), an Eocene cichlid from the Lumbrera formation in Argentina. J. Vert. Paleontol. 30:341–350.

Malabarba, M. C., O. Zuleta, and C. Del Papa. 2006. Proterocara argentina, a new fossil cichlid from the Lumbrera Formation, Eocene of Argentina. J. Vert. Paleontol. 26:267–275.

Marshall, C. R. 2008. A simple method for bracketing absolute divergence times on molecular phylogenies using multiple fossil calibration points. Am. Nat. 171:726–742.

Martin, A. P. and S. R. Palumbi. 1993. Body size, metabolic rate, generation time, and the molecular clock. Proc. Natl. Acad. Sci. USA 90:4087–4091.

McCormack, J. E., J. Heled, K. S. Delaney, A. T. Peterson, and L. L. Knowles. 2011. Calibrating divergence times on species trees versus gene trees: implications for speciation history of Aphelocoma jays. Evolution 65:184–202.

McMahan, C. D., P. Chakrabarty, J. S. Sparks, W. L. Smith, and M. P. Davis. 2013. Temporal patterns of diversification across global cichlid biodiversity (Acanthomorpha: Cichlidae). PLOS ONE 8:e71162.

Meyer, B. S., M. Matschiner, and W. Salzburger. 2015. A tribal level phylogeny of Lake Tanganyika cichlid fishes based on a genomic multi-marker approach. Mol. Phylogenet. Evol. 83C:56–71.

Misof, B., S. Liu, K. Meusemann, R. S. Peters, A. Donath, C. Mayer, P. B. Frandsen, J. Ware, T. Flouri, R. G. Beutel, O. Niehuis, M. Petersen, F. Izquierdo-Carrasco, T. Wappler, J. Rust, A. J. Aberer, U. Aspöck, H. Aspöck, D. Bartel, A. Blanke, S. Berger, A. Böhm, T. R. Buckley, B. Calcott, J. Chen, F. Friedrich, M. Fukui, M. Fujita, C. Greve, P. Grobe, S. Gu, Y. Huang, L. S. Jermiin, A. Y. Kawahara, L. Krogmann, M. Kubiak, R. Lanfear, H. Letsch, Y. Li, Z. Li, J. Li, H. Lu, R. Machida, Y. Mashimo, P. Kapli, D. D. McKenna, G. Meng, Y. Nakagaki, J. L. Navarrete-Heredia, M. Ott, Y. Ou, G. Pass, L. Podsiadlowski, H. Pohl, B. M. von Reumont, K. Schütte, K. Sekiya, S. Shimizu, A. Slipinski, A. Stamatakis, W. Song, X. Su, N. U. Szucsich, M. Tan, X. Tan, M. Tang, J. Tang, G. Timelthaler, S. Tomizuka, M. Trautwein, X. Tong, T. Uchifune, M. G. Walzl, B. M. Wiegmann, J. Wilbrandt, B. Wipfler, T. K. F. Wong, Q. Wu, G. Wu, Y. Xie, S. Yang, Q. Yang, D. K. Yeates, K. Yoshizawa, Q. Zhang, R. Zhang, W. Zhang, Y. Zhang, J. Zhao, C. Zhou, L. Zhou, T. Ziesmann, S. Zou, Y. Li, X. Xu, Y. Zhang, H. Yang, J. Wang, J. Wang, K. M. Kjer, and X. Zhou. 2014. Phylogenomics resolves the timing and pattern of insect evolution. Science 346:763–767.

Murray, A. M. 2000a. Eocene cichlid fishes from Tanzania, East Africa. J. Vert. Paleontol. 20:651–664.

Murray, A. M. 2000b. The Eocene cichlids (Perciformes: Labroidei) of Mahenge, Tanzania. Ph.D. thesis McGill University Montreal.

Murray, A. M. 2001a. The fossil record and biogeography of the Cichlidae (Actinopterygii: Labroidei). Biol. J. Linn. Soc. 74:517–532.

Murray, A. M. 2001b. The oldest fossil cichlids (Teleostei: Perciformes): indication of a 45 million-year-old species flock. Proc. R. Soc. B 268:679–684.

Murrell, B., J. O. Wertheim, S. Moola, T. Weighill, K. Scheffler, and S. L. Kosakovsky Pond. 2012. Detecting individual sites subject to episodic diversifying selection. PLOS Genet. 8:e1002764.

Myers, G. S. 1949. Salt-tolerance of fresh-water fish groups in relation to zoogeographical problems. Bijdragen tot de Dierkunde 28:315–322.

Nabholz, B., S. Glemin, and N. Galtier. 2008. Strong variations of mitochondrial mutation rate across mammals - the longevity hypothesis. Mol. Biol. Evol. 25:120–130.

Near, T. J., A. Dornburg, R. I. Eytan, B. P. Keck, W. L. Smith, K. L. Kuhn, J. A. Moore, S. A. Price, F. T. Burbrink, M. Friedman, and P. C. Wainwright. 2013. Phylogeny and tempo of diversification in the superradiation of spiny-rayed fishes. Proc. Natl. Acad. Sci. USA 110:12738–12743.

O’Reilly, J. E., M. dos Reis, and P. C. J. Donoghue. 2015. Dating tips for divergence-time estimation. Trends Genet. 31:637–650.

Pfeiffer, W. and A. Stamatakis. 2010. Hybrid MPI/Pthreads parallelization of the RAxML phylogenetics code. Ninth IEEE International Workshop on High Performance Computational Biology (HiCOMB 2010), Atlanta Pages 1–8.

Prum, R. O., J. S. Berv, A. Dornburg, D. J. Field, J. P. Townsend, E. M. Lemmon, and A. R. Lemmon. 2015. A comprehensive phylogeny of birds (Aves) using targeted next-generation DNA sequencing. Nature 526:569–573.

Pyron, R. A. 2011. Divergence time estimation using fossils as terminal taxa and the origins of Lissamphibia. Syst. Biol. 60:466–481.

Rabosky, D. L. 2014. Automatic detection of key innovations, rate shifts, and diversity-dependence on phylogenetic trees. PLOS ONE 9:e89543.

Rabosky, D. L., F. Santini, J. Eastman, S. A. Smith, B. Sidlauskas, J. Chang, and M. E. Alfaro. 2013. Rates of speciation and morphological evolution are correlated across the largest vertebrate radiation. Nat. Commun. 4:1958.

Ratnasingham, S. and P. D. N. Hebert. 2007. BOLD: The Barcode of Life Data System (www.barcodinglife.org). Mol. Ecol. Notes 7:355–364.

Ronquist, F. and J. P. Huelsenbeck. 2003. MrBayes 3: Bayesian phylogenetic inference under mixed models. Bioinformatics 19:1572–1574.

Ronquist, F., S. Klopfstein, L. Vilhelmsen, S. Schulmeister, D. L. Murray, and A. P. Rasnitsyn. 2012. A total-evidence approach to dating with fossils, applied to the early radiation of the Hymenoptera. Syst. Biol. 61:973–999.

Roskov, Y., L. Abucay, T. Orell, D. Nicolson, T. Kunze, C. Flann, N. Bailly, P. Kirk, T. Bourgoin, R. E. DeWalt, W. Decock, and A. De Wever. 2015. Species 2000 & ITIS Catalogue of Life. Digital resource at www.catalogueoflife.org/col. Species 2000: Naturalis, Leiden, the Netherlands.

Salzburger, W., B. Van Bocxlaer, and A. S. Cohen. 2014. Ecology and evolution of the African Great Lakes and their faunas. Annu Rev Ecol Evol Syst 45:519–545.

Santini, F., L. J. Harmon, G. Carnevale, and M. E. Alfaro. 2009. Did genome duplication drive the origin of teleosts? A comparative study of diversification in ray-finned fishes. BMC Evol. Biol. 9:194.

Sarich, V. M. and A. C. Wilson. 1967. Immunological time scale for hominid evolution. Science 158:1200–1203.

Schwarzer, J., B. Misof, D. Tautz, and U. K. Schliewen. 2009. The root of the East African cichlid radiations. BMC Evol. Biol. 9:186.

Silvestro, D., N. Salamin, and J. Schnitzler. 2014. PyRate: a new program to estimate speciation and extinction rates from incomplete fossil data. Method Ecol. Evol. 5:1126–1131.

Smith, G. J. D., D. Vijaykrishna, J. Bahl, S. J. Lycett, M. Worobey, O. G. Pybus, S. K. Ma, C. L. Cheung, J. Raghwani, S. Bhatt, J. S. M. Peiris, Y. Guan, and A. Rambaut. 2009. Origins and evolutionary genomics of the 2009 swine-origin H1N1 influenza A epidemic. Nature 459:1122–1125.

Smith, W. L., P. Chakrabarty, and J. S. Sparks. 2008. Phylogeny, taxonomy, and evolution of Neotropical cichlids (Teleostei: Cichlidae: Cichlinae). Cladistics 24:625–641.

Sparks, J. S. and W. L. Smith. 2004. Phylogeny and biogeography of cichlid fishes (Teleostei: Perciformes: Cichlidae). Cladistics 20:501–517.

Sparks, J. S. and W. L. Smith. 2005. Freshwater fishes, dispersal ability, and nonevidence: “Gondwana life rafts” to the rescue. Syst. Biol. 54:158–165.

Stadler, T. 2011. Mammalian phylogeny reveals recent diversification rate shifts. Proc. Natl. Acad. Sci. USA 108:6187–6192.

Stadler, T. and F. Bokma. 2013. Estimating speciation and extinction rates for phylogenies of higher taxa. Syst. Biol. 62:220–230.

Stadler, T., D. Kühnert, S. Bonhoeffer, and A. J. Drummond. 2012. Birth-death skyline plot reveals temporal changes of epidemic spread in HIV and hepatitis C virus (HCV). Proc. Natl. Acad. Sci. USA 110:228–233.

Stamatakis, A. 2006. RAxML-VI-HPC: maximum likelihood-based phylogenetic analyses with thousands of taxa and mixed models. Bioinformatics 22:2688–2690.

Stamatakis, A., P. Hoover, and J. Rougemont. 2008. A rapid bootstrap algorithm for the RAxML web servers. Syst. Biol. 57:758–771.

Starrfelt, J. and L. H. Liow. 2015. How many dinosaur species were there? Fossil bias and true richness estimated using a Poisson sampling model (TRiPS). bioRxiv preprint Pages 1–26.

Stickney, R. R. 1986. Tilapia tolerance of saline waters: a review. Prog. Fish Cult. 48.

Uchida, K., T. Kaneko, H. Miyazaki, S. Hasegawa, and T. Hirano. 2000. Excellent salinity tolerance of Mozambique tilapia (Oreochromis mossambicus): elevated chloride cell activity in the branchial and opercular epithelia of the fish adapted to concentrated seawater. Zool. Sci. 17:149–160.

Vences, M., J. Freyhof, R. Sonnenberg, J. Kosuch, and M. Veith. 2001. Reconciling fossils and molecules: Cenozoic divergence of cichlid fishes and the biogeography of Madagascar. J. Biogeogr. 28:1091–1099.

Vucetich, M. G., M. A. Reguero, M. Bond, A. M. Candela, A. A. Carlini, C. M. Deschamps, J. N. Gelfo, F. J. Goin, G. M. López, E. Ortiz Jaureguizar, R. Pascual, G. J. Scillato-Yané, and E. C. Vieytes. 2007. Mamíferos continentales del Paleógeno argentino: las investigaciones de los últimos cincuenta años. Ameghiniana Publicación Especial 11:239–255.

Wilson, A. C., S. S. Carlson, and T. J. White. 1977. Biochemical evolution. Annu. Rev. Biochem. 46:573–639.

Wright, A. M., G. T. Lloyd, and D. M. Hillis. 2015. Modeling character change heterogeneity in phylogenetic analyses of morphology through the use of priors. Syst. Biol. Advance Access, doi: 10.1093/sysbio/syv122.

Yang, Z. and B. Rannala. 2006. Bayesian estimation of species divergence times under a molecular clock using multiple fossil calibrations with soft bounds. Mol. Biol. Evol. 23:212–226.

Zhang, C., T. Stadler, S. Klopfstein, T. A. Heath, and F. Ronquist. 2015. Total-evidence dating under the fossilized birth-death process. Syst. Biol. Advance Access, doi: 10.1093/sysbio/syv080.

Zuckerkandl, E. and L. Pauling. 1962. Molecular disease, evolution, and genic heterogeneity. Pages 189–225 in Horizons in Biochemistry (M. Kasha and B. Pullman, eds.). Academic Press, New York.

